# RNA Sensing and Innate Immunity Constitutes a Barrier for Interspecies Chimerism

**DOI:** 10.1101/2023.03.07.531624

**Authors:** Yingying Hu, Hai-Xi Sun, Masahiro Sakurai, Amanda E. Jones, Lizhong Liu, Tianlei Cheng, Canbin Zheng, Jie Li, Benjamin Ravaux, Zhou Luo, Yi Ding, Tianbin Liu, Yan Wu, Elizabeth H. Chen, Zhijian J. Chen, John M. Abrams, Ying Gu, Jun Wu

## Abstract

Interspecies chimera formation with human pluripotent stem cells (PSCs) holds great promise to generate humanized animal models and provide donor organs for transplant. However, the approach is currently limited by low levels of human cells ultimately represented in chimeric embryos. Different strategies have been developed to improve chimerism by genetically editing donor human PSCs. To date, however, it remains unexplored if human chimerism can be enhanced in animals through modifying the host embryos. Leveraging the interspecies PSC competition model, here we discovered retinoic acid-inducible gene I (RIG-I)-like receptor (RLR) signaling, an RNA sensor, in “winner” cells plays an important role in the competitive interactions between co-cultured mouse and human PSCs. We found that genetic inactivation of *Ddx58/Ifih1-Mavs-Irf7* axis compromised the “winner” status of mouse PSCs and their ability to outcompete PSCs from evolutionarily distant species during co-culture. Furthermore, by using *Mavs*-deficient mouse embryos we substantially improved unmodified donor human cell survival. Comparative transcriptome analyses based on species-specific sequences suggest contact-dependent human-to-mouse transfer of RNAs likely plays a part in mediating the cross-species interactions. Taken together, these findings establish a previously unrecognized role of RNA sensing and innate immunity in “winner” cells during cell competition and provides a proof-of-concept for modifying host embryos, rather than donor PSCs, to enhance interspecies chimerism.

## Main text

The technique of generating interspecies chimeras using human pluripotent stem cells (hPSCs) is a promising *in vivo* platform to study human development and offers a potential source for growing human donor organs in animals^1, 2^. Although robust chimerism can be achieved between closely related species^3, 4^, it is far more difficult to generate chimeras between evolutionarily distant species^5^. The low chimerism of human cells in animals (e.g., mice and pigs) is presumably due to multiple xenogeneic barriers during early development, which include but are not limited to differences in developmental pace, incompatibility in cell adhesion molecules, and interspecies cell competition. Several strategies have been developed to improve human cell chimerism in animal embryos by genetically inhibiting human cell apoptosis^6–10^. However, these strategies are not practical for future use in regenerative medicine as the modified genes and pathways are mostly oncogenic. Improving survival and chimerism of unmodified donor hPSCs through editing the host embryos will be a preferred solution but has not been explored.

We previously developed an interspecies PSC co-culture system and discovered a competitive interaction between primed but not naïve human and mouse PSCs, whereby loser hPSCs are eliminated by winner mouse epiblast stem cells (mEpiSCs) through apoptosis. Genetic inactivation of *MyD88*, *P65* or *P53* in hPSCs could overcome human-mouse PSC competition, thereby improving human cell survival and chimerism in early mouse embryos^10^. In contrast to loser hPSCs, however, little is known regarding what enacts the winner status of mEpiSCs. To this end, we performed RNA sequencing (RNA-seq) of separately cultured and co-cultured mEpiSCs. H9 human embryonic stem (ES) cells and mEpiSCs were labeled with enhanced green fluorescent protein (eGFP) and monomeric Kusabira Orange (mKO), respectively. We isolated mEpiSCs from day 1-3 co-cultures and separated cultures by fluorescence-activated single cell sorting (FACS) for RNA-seq (**Fig. 1a**, **Extended Data Fig. 6a, b**). Comparative transcriptome analysis identified 422, 301 and 77 upregulated genes on days 1, 2 and 3, respectively, in co-cultured versus separately cultured mEpiSCs (co-culture upregulated genes, or co-URGs) (**Fig. 1b**). Kyoto Encyclopedia of Genes and Genomes (KEGG) pathway analyses were performed using these co-URGs. Interestingly, many overrepresented pathways in days 1 and 2 co-URG groups were related to virus infection and innate immunity (**Extended Data Fig. 1a-c, Supplementary Table 1**). Among them, retinoic acid-inducible gene I (RIG-I)-like receptor (RLR) signaling pathway represents one of the major RNA sensing systems in mammalian cells, which can be activated by viral and host-derived RNAs^11, 12^. RLR activation results in the transcriptional induction of type I interferons that mounts an antiviral host response^13^. RLR signaling participates in the maintenance of homeostatic genetic stability of cell populations and self-protection^11, 14^. Several key component genes of the RLR pathway were significantly upregulated in days 1-2 co-cultured versus separately cultured mEpiSCs, including the cytoplasmic RNA sensors *Ddx58* (RIG-I)^15^ and *Ifih1* (MDA5)^16, 17^ and the downstream transcription factor *Irf7*^18^ (**Fig. 1c, d**, **Extended Data Fig. 1d-f**).

**Fig. 1.**
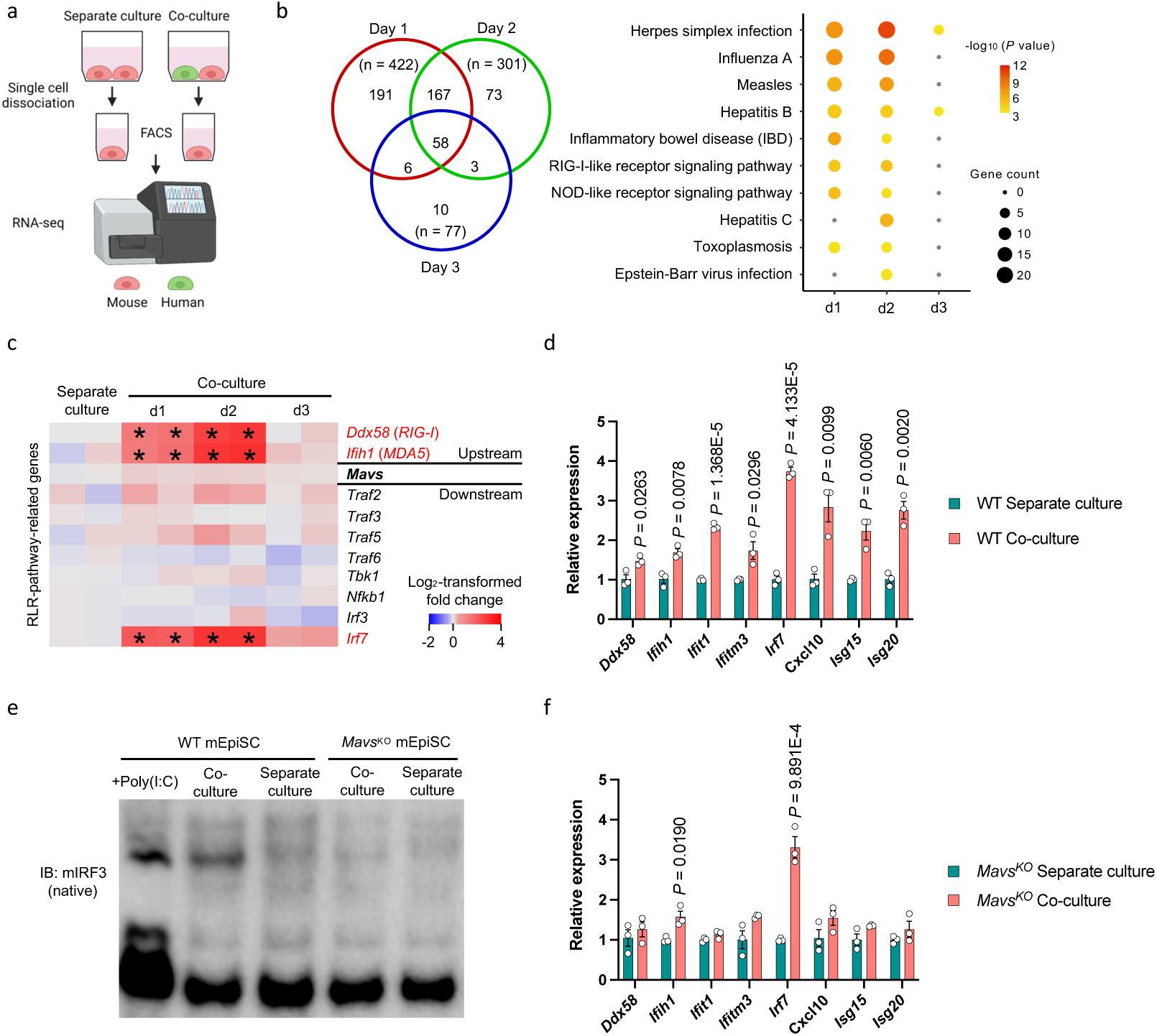
| RLR signaling activation in mEpiSCs co-cultured with H9 hES cells. **a,** Schematic of human and mouse PSC co-culture and RNA-seq experimental setup. **b,** Left, Venn diagram showing the numbers of co-URGs in mEpiSCs. Right, KEGG pathway analysis of co-URGs in mEpiSCs. The color of each dot represents log_10_-transformed *P* value of each term, while the dot size represents the number of co-URGs in each term. Terms that are not significantly enriched are shown in gray. **c,** Heatmap showing fold changes of RLR-pathway-related genes in co-cultured versus separately cultured mEpiSCs. Blue and red represent log_2_-transformed fold changes < 0 and > 0, respectively. Asterisks indicate statistically differentially expressed. **d,** RT-qPCR analysis of relative expression levels of RLR-pathway-related genes in co-cultured WT mEpiSCs compared to separately cultured WT mEpiSCs. n = 3, biological replicates. Data are mean ± s.e.m. *P* values determined by unpaired two-tailed t-test. **e,** Native gel electrophoresis detecting IRF3 dimerization in WT mEpiSCs transfected with poly (I:C), co-cultured and separately cultured WT mEpiSCs, co-cultured and separately cultured *Mavs*^KO^ mEpiSCs. **f,** RT-qPCR analysis of relative expression levels of RLR-pathway-related genes in co-cultured *Mavs*^KO^ mEpiSCs compared to separately cultured *Mavs*^KO^ mEpiSCs. n = 3, biological replicates. Data are mean ± s.e.m. *P* values determined by unpaired two-tailed t-test.

Next, we determined whether the RLR pathway is intact in mEpiSCs. To this end, we transfected mEpiSCs with polyinosinic:polycytidylic acid (also known as poly (I:C)), a double-stranded RNA analogue that can activate RLRs^19, 20^. Native gel electrophoresis and western blotting revealed IRF3 dimerization, demonstrating an activated RLR signaling in poly (I:C) transfected mEpiSCs (**Fig. 1e**). To confirm RLR pathway activation in mEpiSCs co-cultured with H9 hES cells, FACS-isolated mEpiSCs from day 2 co-culture and separate culture were subjected to protein extraction and native gel electrophoresis. Consistent with RNA-seq results, we found RLR pathway was activated as evidenced by IRF3 dimerization in co-cultured but not separately cultured mEpiSCs (**Fig. 1e**). RLR pathway activation in co-cultured mEpiSCs seems to be contact-dependent as we did not observe upregulation of RLR-pathway-related genes in mEpiSCs co-cultured with H9 hES cells in trans-wells (**Extended Data Fig. 1g**). Interestingly, when *P65^KO^*-hiPSCs were used for co-culture, upregulated expression of RLR-pathway-related genes were still observed in mEpiSCs (**Extended Data Fig. 1h, i**), demonstrating RLR signaling activation in mEpiSCs is independent of the outcomes of hPSCs during co-culture. In contrast to mEpiSCs, there was no apparent activation of the RLR signaling pathway in co-cultured H9 hES cells (**Extended Data Fig. 2a-i, Supplementary Table 2)**. Collectively, during co-culture with loser hPSCs, our results uncovered an intriguing cell contact-dependent activation of RLRs in winner mEpiSCs.

Mitochondrial anti-viral signaling protein (MAVS) is the central hub for RLR sensors and downstream signaling^18, 21–23^. To determine whether RLR signaling contributes to human-mouse PSC competition, we knocked out *Mavs* in mEpiSCs (*Mavs*^KO^ mEpiSCs) and confirmed *Mavs* deficiency did not affect self-renewal, proliferation and expression of key pluripotency transcription factors (**Extended Data Fig. 3a-c**). IRF3 dimerization signal was lost in day 2 co-cultured *Mavs*^KO^ mEpiSCs (**Fig. 1e**) and most RLR-pathway-related genes were expressed at comparable levels to separate cultures (**Fig. 1f**), demonstrating *Mavs* knockout is effective in blocking RLR signaling in co-cultured mEpiSCs. Time-lapse confocal microscopy revealed better survival of H9 hES cells during co-culture with *Mavs*^KO^ mEpiSCs than with WT mEpiSCs (**Supplementary Video 1**). Next, we calculated the cell densities (cell number per cm^2^) of live H9 hES cells and mEpiSCs in co-culture and separate culture daily until they grew to confluency (days 1-5). A more rapid elimination of H9 hES cells was observed during co-culture with WT than with *Mavs*^KO^ mEpiSCs (**Fig. 2a**). These results suggest compromised competitiveness of *Mavs*^KO^ mEpiSCs. We also tested higher cell plating densities. Notably, on day 3, significantly more H9 hES cells were found in co-cultures with *Mavs*^KO^ than with WT mEpiSCs (**Fig. 2b**, **Extended Data Fig. 3d**). To confirm this finding with an independent method, we performed automatic cell recognition and unbiased counting by using an image analysis software. To this end, eGFP-H2B labeled WT or *Mavs*^KO^ mEpiSCs and/or mScarlet-H2B labeled H9 hES cells were plated in micropatterned plates. Fluorescence images were recorded over time for each micropatterned colony using confocal microscopy followed by image analysis and automated cell counting (see Methods). In consistent, on day 3, significantly higher numbers of live H9 hES cells were present in co-cultures with *Mavs*^KO^ than with WT mEpiSCs (**Extended Data Fig. 3e, f**).

**Fig. 2.**
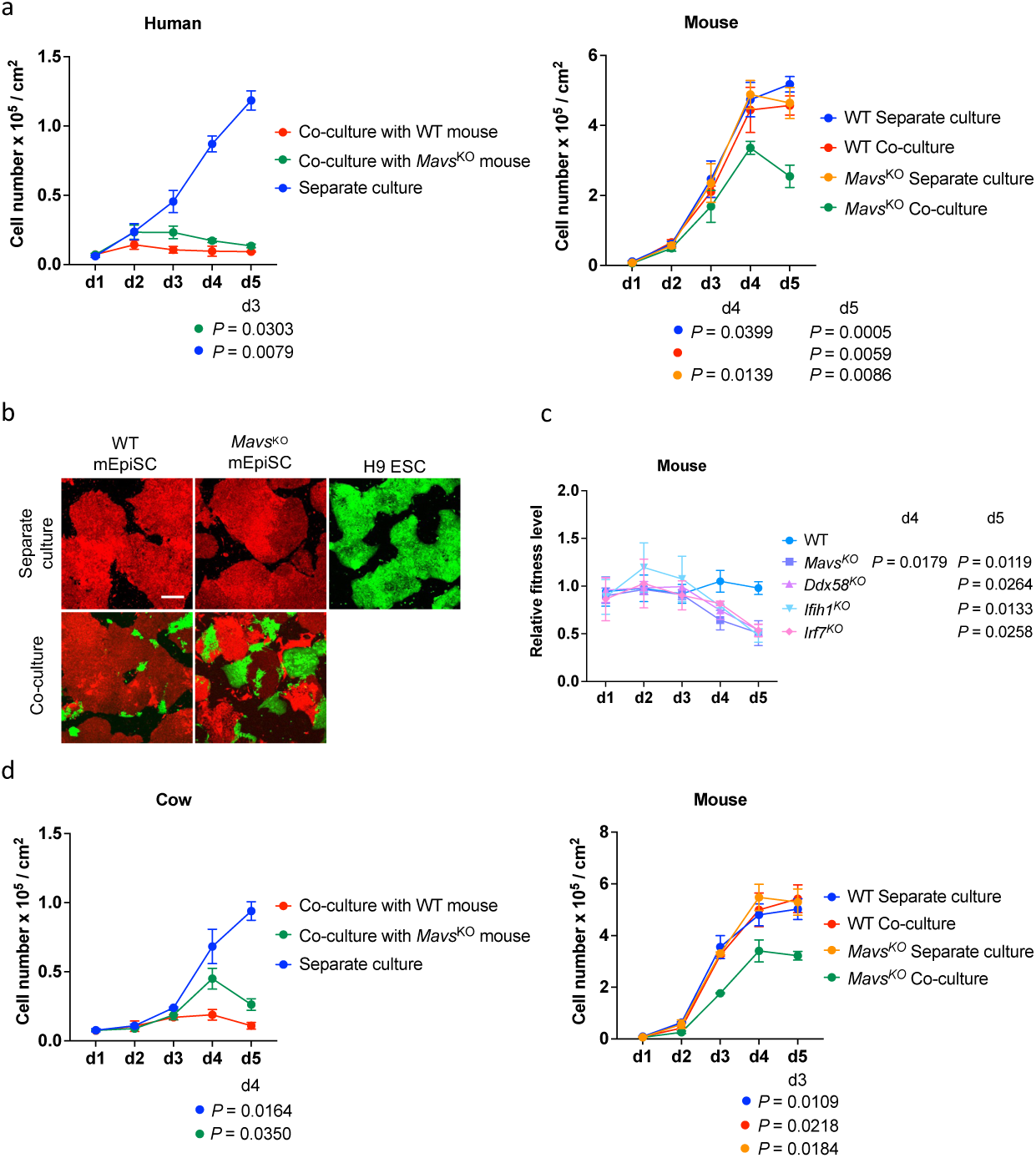
| Compromised cell competitiveness and viability of *Mavs-deficient* mEpiSCs. **a,** Left, growth curves of H9 hES cells in co-culture with WT mEpiSCs (red), co-culture with *Mavs*^KO^ mEpiSCs (green) and separate culture (blue). Right, growth curves of WT mEpiSCs in co-culture (red) with H9 hES cells and separate culture (blue), *Mavs*^KO^ mEpiSCs in co-culture (green) with H9 hES cells and separate culture (orange). n = 6, biological replicates. **b,** Representative fluorescence images of day-3 co-cultured and separately cultured WT mEpiSCs (red), *Mavs*^KO^ mEpiSCs (red) and H9 hES cells (green). Scale bar, 200 μm. **c,** Growth curves of co-cultured WT, *Mavs*^KO^, *Ddx58*^KO^, *Ifih1*^KO^ and *Irf7*^KO^ mEpiSCs normalized to corresponding separate cultures. n = 6 (WT), n = 6 (*Mavs*^KO^), n = 3 (*Ddx58*^KO^), n = 3 (*Ifih1*^KO^) and n = 3 (*Irf7*^KO^) independent culture experiments. **d,** Left, growth curves of bovine ES cells in co-culture with WT mEpiSCs (red), co-culture with *Mavs*^KO^ mEpiSCs (green) and separate culture (blue). Right, growth curves of WT mEpiSCs in co-culture (red) with bovine ES cells and separate culture (blue), *Mavs*^KO^ mEpiSCs in co-culture (green) with bovine ES cells and separate culture (orange). n =3, biological replicates. All data are mean ± s.e.m. *P* values determined by one-way ANOVA with Dunnett’s multiple comparison.

Interestingly, during co-culture with H9 hES cells, more *Mavs*^KO^ mEpiSCs appeared to undergo cell death than WT mEpiSCs (**Supplementary Video 1**). In addition, starting from day 4, in contrast to WT mEpiSCs, significantly lower densities of *Mavs*^KO^ mEpiSCs were found in co-cultures than separate cultures (**Fig. 2a**). These results suggest reduced viability of *Mavs*^KO^ mEpiSCs during co-culture with H9 hES cells. In addition to *Mavs*, we individually knocked out genes upstream (*Ddx58*, *Ifih1)* and downstream (*Irf7) of Mavs* in the RLR pathway in mEpiSCs. *Ddx58*^KO^-, *Ifih1*^KO^-and *Irf7*^KO^-mEpiSCs all could be passaged long term in culture and maintained the expression of OCT4 and SOX2 (**Extended Data Fig. 4a, b)**. Similar to *Mavs*, deficiency of *Ddx58*, *Ifih1* and *Irf7* also led to the significant reduction in mEpiSC densities during co-cultures with H9 hES cells (**Fig. 2c**, **Extended Data Fig. 4c**). Together, these results reveal that both cell competitiveness and viability were compromised in winner mEpiSCs by depleting *Mavs*, and demonstrate the important role of RLR pathway in human-mouse PSC competition.

In addition to human cells, mEpiSCs could outcompete rhesus macaque and bovine ES cells during co-cultures^10^. Next, we studied whether RLR signaling was involved in PSC competition between mouse and other species. We co-cultured WT and *Mavs*^KO^ mEpiSCs with rhesus (ORMES23) and bovine ES cells, respectively. On days 4-5, there were evidently more rhesus and bovine ES cells remained in co-cultures with *Mavs*^KO^ than with WT mEpiSCs, while significantly less *Mavs*^KO^ mEpiSCs were found in co-cultures than WT mEpiSCs (**Fig. 2d**, **Extended Data Fig. 5a-c, Supplementary Video 2)**. These findings demonstrate that, in the presence of rhesus and bovine ES cells, *Mavs* deficiency also diminished the competitiveness and fitness of mEpiSCs. When compared with separately cultured mEpiSCs, many RLR-pathway-related genes were upregulated in days 1-3 mEpiSCs co-cultured with bovine ES cells (**Extended Data Fig. 5d**), consistent with what we found in human-mouse PSC co-cultures. Collectively, these results suggest conserved roles of RLR signaling during competitive interactions of mEpiSCs with primed PSCs from other species.

To determine whether the suppression of RLR signaling in host mouse embryos could improve donor hPSC survival, we microinjected the same number (10) of eGFP labeled hiPSCs into WT and *Mavs*^-/-^ mouse^24^ blastocysts followed by ex vivo culture^25^ (**Fig. 3a**). After 3 and 5 days, eGFP signals were detected in most *Mavs*^-/-^ embryos injected with hiPSCs, whereas fewer WT embryos contained eGFP+ human cells (**Fig. 3b**, **c)**. To study human cell survival in *Mavs*-deficient mouse embryos in vivo, we electroporated WT mouse embryos with Cas9 protein and *Mavs* targeting sgRNAs at zygote stage, microinjected eGFP+ hiPSCs to the blastocysts and then performed embryo transfer (**Fig. 3a**). We found the percentages of embryonic day (E) 8.5-9.5 mouse embryos containing eGFP+ signals were 18.75% (6 out of 32) for *Mavs*^-/-^, but only 5.26% (1 out of 19) for *Mavs*^+/-^ and 0% (0 out of 9) for *Mavs*^+/+^ (**Fig. 3d**). The presence of human cells in E8.5-9.5 *Mavs*^-/-^ embryos was validated by immunostaining of eGFP (**Fig. 3e**) and genomic PCR using human-specific *Al u* (*TPA25-Alu*) primers^26^ (**Fig. 3f**). Together, these results demonstrate genetic inactivation of RLR signaling in the host could improve unmodified donor human cell survival in early mouse embryos.

**Fig. 3.**
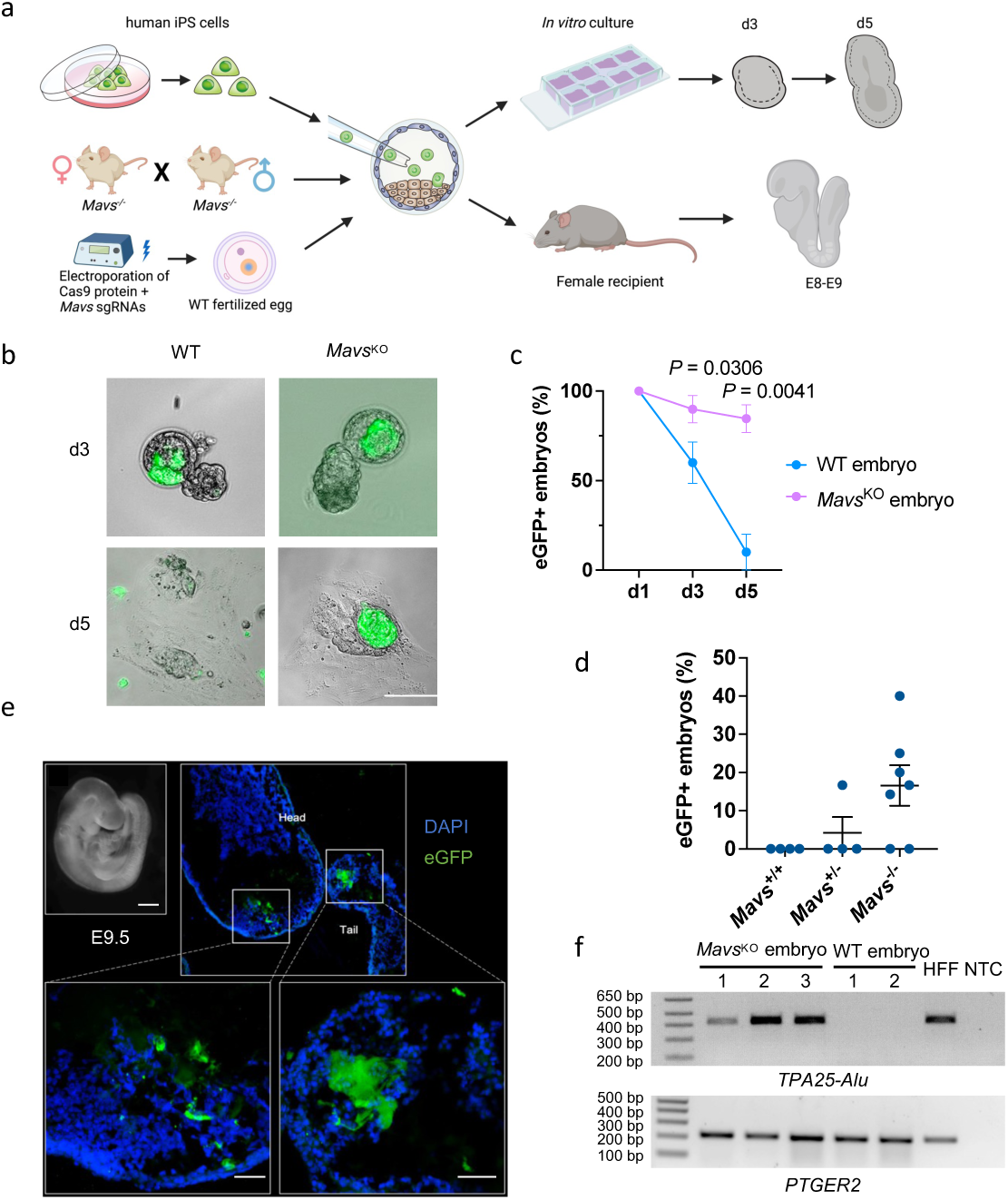
| Genetic inactivation of *Mavs* in mouse embryos improves donor human cell survival during chimera formation. **a,** Schematic showing the generation of ex vivo and in vivo human-mouse chimera using *Mavs*-deficient mouse blastocysts. **b,** Representative brightfield and fluorescence merged images of WT and *Mavs*^KO^ mouse embryos cultured for 3 days (d3) and 5 days (d5) after blastocyst injection with hiPSCs. Scale bar, 100 μm. **c,** Line graphs showing the percentages of eGFP+ mouse embryos at indicated time points during ex vivo culture after injecting hiPSCs into WT and *Mavs*^KO^ blastocysts. n = 3 (WT) and n = 3 (*Mavs*^KO^) independent injection experiments. Data are mean ± s.e.m. *P* values determined by unpaired two-tailed t-test. **d,** Dot plot showing the percentages of eGFP+ E8.5–9.5 mouse embryos derived from injecting hiPSCs into WT and *Mavs*KO blastocysts. Each blue dot represents one embryo transfer experiment. n = 4 (*Mavs*^+/+^), n = 4 (*Mavs*^+/-^) and n = 7 (*Mavs*^-/-^), independent experiments. **e,** Representative immunofluorescence images showing contribution of eGFP-labelled hiPSCs in E9.5 mouse embryos. Embryo sections were stained with antibody against eGFP and DAPI. Scale bars, 500 μm (whole embryo) and 50 μm (insets). **f,** Genomic PCR analysis of E8.5-9.5 mouse embryos derived from injecting hiPSCs into WT and *Mavs*^KO^ blastocysts. *TPA25-Alu* denotes a human-specific primer; *PTGER2* was used as a loading control. HFF, HFF-hiPS cells. NTC, non-template control. This experiment was repeated independently three times with similar results.

Horizontal RNA transfer mediated by exosomes, free extracellular ribonucleoprotein (RNP) particles, or tunneling nanotubes has been observed in mammals^27–30^. We speculated that the observed RLR signaling pathway activation in mEpiSCs during co-culture with hPSCs was likely caused by interspecies (human-to-mouse) horizontal RNA transfer. To look for evidence, we revisited our RNA-seq (bulk) data, which was generated from FACS-sorted cells based on fluorescence (human, eGFP^10^; mouse, mKO) to minimize cross-species contamination (**Fig. 1a**). To find out whether human RNAs are present in mEpiSCs, we first identified mouse-and human-specific reads, and used only these species-specific reads to calculate human genome mapping rates for each sequenced sample (**Fig. 4a**). Interestingly, we detected a small proportion of human-specific reads in co-cultured mEpiSCs (0.57%, 1.30% and 0.33%, for day 1, day 2, and day 3, respectively), which was most prominent on day 2 and consistent with our finding on RLR signaling activation (**Fig. 4b, 1b, 1c**). Similar results were obtained from mEpiSCs co-cultured with *P65^KO^*-hiPSCs (**Extended Data Fig. 6c**). We also performed the same analysis using RNA-seq data generated from non-contact (transwell) co-cultures (**Extended Data Fig. 6d**). Interestingly, human-specific reads were barely detectable in both transwell co-cultured (0.05%) or separately cultured mEpiSCs (0.03%) (**Fig. 4b**ll. Together, these results reveal the presence of human transcripts in co-cultured mEpiSCs, and suggest putative horizontal RNA transfer through physical connections (e.g., membrane nanotubesll^28^llll (e.g., extracellular vesiclesll^27^.

**Fig. 4.**
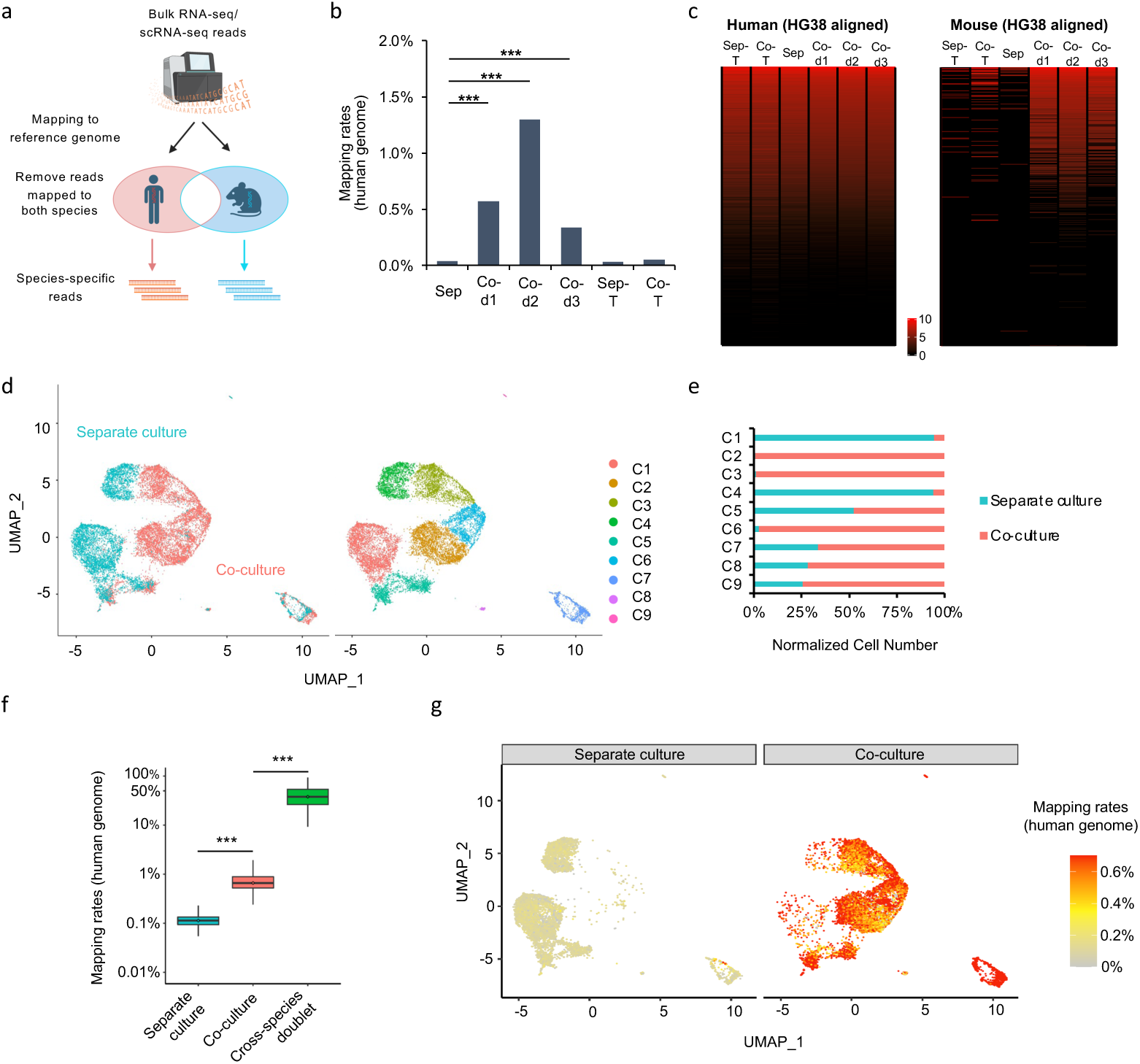
| Presence of human transcripts in co-cultured mEpiSCs revealed by RNA sequencing. **a,** Computational pipeline to identify species-specific bulk/single-cell RNA-seq reads. See Methods for details. **b,** Mapping rates of human genome in mEpiSCs determined by bulk RNA-seq. Asterisks indicate statistically significant differences: (***) *P* < 0.001; Fisher’s exact test. **c,** Heatmap showing global expression profiles of human genes expressed in H9 hES cells and detected in mEpiSCs. Genes are sorted by expression levels from high to low in H9 hES cells. Log_2_-transformed expression levels were used to draw this plot. For **b** and **c**, Sep, Co-d1, Co-d2, Co-d3, Sep-T and Co-T represent separate culture, co-culture on days 1, 2, 3, separate culture in transwell and co-culture in transwell, respectively. **d,** Clustering of separately cultured and co-cultured mEpiSCs by scRNA-seq analysis. Each point represents one single cell, colored according to cluster. **e,** Percentage of separately cultured and co-cultured mEpiSCs in each cluster. **f,** Mapping rates of human genome in mEpiSCs determined by scRNA-seq. Asterisks indicate statistically significant differences: (***) *P* < 0.001; Wilcoxon test. Cross-species doublets served as a negative control. **g,** UMAP visualization showing mapping rates of human genome in separately cultured (left) and co-cultured (right) mEpiSCs.

We then determined the abundance of human transcripts in co-cultured mEpiSCs and found it was positively correlated with gene expression levels in separately cultured H9 hES cells (**Fig. 4c**). Stress-induced transcription of retrotransposable elements (REs) has recently been recognized to play an important role in RNA immune response^31–34^. Therefore, we interrogated multi-mapped reads in these RNA-seq datasets using pipelines designed^35^. Intriguingly, we detected the presence of human RNAs coding for both young and ancient retroelements in mouse EpiSCs, which was contact-dependent and correlated with abundance and peaked at day 2 (**Extended Data Fig. 7a, b**). Of note is that the expression levels of human retroelements in H9 hES cells were not affected by culture conditions (**Extended Data Fig. 7c-f**). Therefore, retroelement RNAs may contribute to RLR signaling activation in mouse cells. In addition, the non-functional exogenous eGFP sequences could also be detected in mEpiSCs specifically under co-cultured condition (**Extended Data Fig. 6e**), providing further evidence that the putative interspecies horizontal RNA transfer process is random instead of selectively active transport.

To rule out the possibility that the co-cultured mEpiSCs collected by FACS was contaminated with small amounts of human cells, we performed single cell RNA-seq (scRNA-seq), using mEpiSCs isolated from day 2 co-culture and separate culture. After filtering out cells with low quality, we obtained transcriptome datasets from 16,225 cells including 6,969 and 9,256 cells in separate and co-cultured conditions, respectively (**Fig. 4d**, **Extended Data Fig. 8a**). We performed unsupervised clustering and identified 9 clusters (**Fig. 4d**). We observed significant differences in cell composition between separate and co-cultures: C2, C3 and C6 clusters predominantly consisted of co-cultured cells, whereas C1 and C4 clusters were mainly from cells in separate culture (**Fig. 4e**). To account for possible cell fusion or doublet generated in scRNA-seq library preparation process, as a control, we included “cross-species doublets” identified by multi-genome analysis by Cell Ranger. Consistent with bulk RNA-seq, scRNA-seq data also revealed significantly elevated mapping rates (∼1.2%) to the human genome in co-cultured mEpiSCs, which was significantly higher than in separately cultured mEpiSCs (∼0.1%) and much lower than “cross-species doublet” (∼40.2%) (**Fig. 4f**). UMAP visualization revealed that the presence of human RNAs could be detected in many co-cultured mEpiSCs (**Fig. 4g**). In addition, eGFP transcripts from human cells were significantly increased in co-cultured versus separately cultured mEpiSCs, which was also significantly lower than those in “cross-species doublet” (**Extended Data Fig. 8b-d**). Next, based on the scRNA-seq data, we performed expression correlation analysis of human transcripts, and found high Pearson correlation coefficients (PCCs) between co-cultured mEpiSCs and H9 hES cells, especially, genes with higher expression level in H9 hES cells are more likely to be detected in co-cultured mEpiSCs (**Extended Data Fig. 8e**). These results confirmed the presence of human RNA in co-cultured mEpiSCs at single cell level and further suggest that the putative human-to-mouse RNA transfer is non-selective.

In summary, we discovered RLR signaling pathway was activated in mEpiSCs during interspecies primed PSC competition. Genetic perturbation of key players in the RLR signaling pathway compromised the “winner” status of mEpiSCs, and thereby resulting in better survival of “loser” PSCs from evolutionarily distant species, including humans, rhesus monkeys and cattle. We further showed that by using *Mavs*-deficient mouse embryos as hosts, survival of unmodified donor hPSCs could be improved. In addition, we detected the presence of human transcripts encoding unique genes and retrotransposable elements in co-cultured mEpiSCs, putatively as a result a non-selective and contact dependent human-to-mouse RNA transfer. Future studies are warrantied to dissect the detailed molecular mechanisms and functional consequences underlying this horizontal RNA transfer between PSCs from different species.

Our study uncovers an unexpected and intriguing role of RNA innate immunity in interspecies PSC competition and chimerism, and it’ll be interesting to determine whether the same mechanism also operates in “winner” cells in classical cell competition models such as *Minute* mutant^36^ and *dMyc*^37^. By suppressing the RLR pathway in mouse embryos, we demonstrate the feasibility of modifying the host embryos to boost interspecies chimerism of donor PSCs, providing a framework for generating human tissues in animals that are more suitable for transplant in the future.

## Acknowledgements

We thank T. Hishida (Wakayama Medical University, Japan) for providing pCAG-IP-mKO and pCAG-IP-eGFP plasmids; Aryeh Warmflash lab for providing piggyBac-eGFP-H2B and piggyBac transposase plasmids; B. Li and J. Ye for generating single cell RNA sequencing library; L. Zhang and Y. Zhao for technical support; the services provided by China National GeneBank and NCBI Gene Expression Omnibus. J.W. is a New York Stem Cell Foundation (NYSCF)– Robertson Investigator and Virginia Murchison Linthicum Scholar in Medical Research and this work is funded by CPRIT (RR170076), NYSCF, NIH (HD103627-01A1, not used for human-mouse chimera work), and Welch (854671). E.H.C. is supported by NIH grants R01AR075005 and R35GM136316. Y.H., T.L. and Y.G. is supported by Guangdong Provincial Key Laboratory of Genome Read and Write (no. 2017B030301011). We thank BioHPC at UTSW for providing high performance computational resources. J.M.A.is supported by NIH grants R01GM115682 and R01CA222579.

## Author contributions

J.W. conceptualized the idea, provided research support, designed, analyzed, and interpreted the results, and wrote the manuscript. Y.H. contributed to the study design, generated all the KO cell lines, performed most of the cell competition experiments, and wrote the manuscript. H.S., J.L., Y.W. and Y.G. performed bulk RNA sequencing and single-cell RNA sequencing analysis. Y.G. also contributed to the study design and co-supervised the project. M.S. performed blastocyst microinjections and embryo transfers. A.E.J. and J.M.A conceived, designed and performed bioinformatic analyses focusing on the retrotransposable elements. L.L. performed micropattern co-cultures and imaging assay. T.C. and Z.J.C. performed western blotting and native gel electrophoresis. C.Z. contributed to the study design and preparation of bulk RNA sequencing samples. B.R., Z.L. and E.H.C. contributed to time-lapse imaging experiments using micropatterned coverslips. Y.D. helped with mouse zygotes electroporation and blastocysts collection. T.L., C.Z., A.E.J., and J.M.A helped write the manuscript. Y.H., H.S. and Z.L. organized the figures. All authors reviewed the manuscript.

## Methods

### Animals and ethical review

CD-1 (ICR), C57BL/6 and BDF1 mice were purchased from Envigo (Harlen), Charles River, or Jackson Laboratory. Homozygous *Mavs* knockout mice were kindly gifted from Prof. Zhijian J. Chen^24^. Mice were housed in 12-hr light/12-hr dark cycle at 22.1–22.3°C and 33–44% humidity. All procedures related to animals were performed in accordance with the ethical guidelines of the University of Texas Southwestern Medical Center (UTSW). Animal protocols were reviewed and approved by the UTSW Institutional Animal Care and Use Committee (IACUC) before any experiments were performed (Protocols #2018-102430). All experiments followed the 2021 Guidelines for Stem Cell Research and Clinical Translation released by the International Society for Stem Cell Research (ISSCR). All human–mouse ex vivo and in vivo interspecies chimeric experimental studies were reviewed and approved by UT Southwestern Stem Cell Oversight Committee (SCRO) (registration 14).

### Primed PSC culture

Human ES cell line H9 (WA09) were obtained from WiCell and authenticated by short tandem repeat (STR) profiling. HFF-hiPS cells, mouse EpiSCs, rhesus macaque ES cells, and bovine ES cells were generated as previously described^5, 38, 39^. Human primed PSCs were either cultured on plates coated with Matrigel (BD Biosciences) in mTeSR1 medium (STEMCell Technologies) or on MEFs in NBFR medium, which contains DMEM/F12 (Invitrogen) and Neurobasal medium (Invitrogen) mixed at 1:1 ratio, 0.5× N2 supplement (Invitrogen), 0.5× B27 supplement (Invitrogen), 2 mM GlutaMax (Gibco), 1× nonessential amino acids (NEAA, Gibco), 0.1 mM 2-mercaptoethanol (Sigma-Aldrich), 20 ng mL^−1^ FGF2 (PeproTech), 2.5 μM IWR1 (Sigma-Aldrich), and 1 mg mL^−1^ BSA (low fatty acid, MP Biomedicals). Mouse, rhesus and bovine primed PSCs were all cultured on MEFs in NBFR medium. Primed PSCs cultured in NBFR medium were passaged using TrypLE (human, rhesus and bovine) at 1:10 split ratio every 4–5 days, and 1:30 split ratio (mouse) every 3–4 days. Human primed PSCs cultured in mTeSR1 medium on Matrigel were passaged every five days using Dispase (STEMCell Technologies) at 1:10 split ratio.

### Generation of fluorescently labelled PSCs

We used pCAG-IP-mKO and pCAG-IP-eGFP to label PSCs as previously described^10^. For generating nucleus labelled PSCs, we used piggyBac transposon vectors expressing eGFP-H2B and mScarlet-H2B. In brief, 3 μg piggyBac-eGFP-H2B or piggyBac-mScarlet-H2B plasmids were transfected together with 1 μg piggyBac transposase plasmid into 1 × 10^6^–2 × 10^6^ dissociated PSCs using NEPA21 electroporator (Nepa Gene) following the protocol recommended by the manufacturer. eGFP^+^ or mScarlet+ cells were collected by FACS at 48 h after transfection and replated. Single clones with stable fluorescence were manually picked between 7 and 14 days and further expanded clonally.

### Interspecies PSC co-culture

PSCs from different species were seeded onto MEF-coated plates and either cultured separately or mixed at different ratios for co-cultures in NBFR medium. The seeding ratio and density were empirically tested and decided on the basis of cell growth rate. For 5-days-co-culture assays between human-mouse, rhesus-mouse and bovine-mouse PSCs, cells were seeded at a 4:1 ratio at a density of 1.25 × 10^4^ cells cm^−2^. For human-mouse 3-days-co-culture, seeding density was increased to 1.875 × 10^4^ cells cm^−2^ but seeding ratio was kept at 4:1. For transwell co-culture experiments, Millipore Transwell 0.4 μm PET hanging inserts (Millicell, MCH12H48) were placed into 12-well plates. Then coverslips were placed into both the upper insert and the bottom well, mEpiSCs and hES cells were seeded on the top insert and bottom well, respectively. At each of the indicated time points, cell concentration was manually counted and calculated, and the percentages of each cell line were determined using FACSCalibur system (BD Bioscience). Total cell numbers (tN) for each species in co-cultures or separate cultures were determined by multiplying total cell volume (V) with cell concentration (CC) and percentage (P). tN = V × CC × P. Cell density (cells cm^−2^) was calculated by dividing the total cell number by the surface area.

### Micropattern imaging and automated cell counting

The micropattern-based imaging was performed on CYTOO micropattern plates (CYTOOplate 96w Custom A700-P400 A). Specifically, the micropattern wells were coated with 0.5% (v/v) Matrigel at 4 °C, O/N, and MEF feeder cells were seeded in each well one day before the experiment starts. On the day the experiment starts (day 0), unattached MEF feeder cells were washed out and then eGFP-H2B labeled WT or *Mavs*^KO^ mEpiSCs and/or mScarlet-H2B labeled H9 hES cells were seeded into each well separately or mixed as indicated in the main text. On days 1-3, individual micropattern colonies were tracked by live imaging. Cells were then fixed and DAPI stained on day 3. Fluorescence imaging was performed on Nikon CSU-W1 SoRa spinning disk confocal microscope with 20x (0.75 NA, WD1 (air)) objective. Z-stacks were acquired for each micropattern colony with 1µm step, approximately 30-75µm range in Z was covered depending on the thickness of each colony. Imaging experiments were repeated at least three times with similar results. Cell counting was performed on Imaris (version 9.1, Oxford Instruments) by using the SPOTS function. Same parameters for computing of human and mouse cells were applied.

### Time-lapse imaging and analysis

For time-lapse imaging of co-cultures using bowtie-shaped microwells^40^, the photoresist template was fabricated by negative photolithography as previously described^41^. The chrome mask was manufactured by the University of Texas at Dallas, and KMPR 1050 photoresist (Microchem, Westborough, MA) was used following the manufacturers’ instructions. Silicon (PDMS) molds were fabricated from Sylgard 184 Silicon Elastomer (#24236-10 – Electron Micro Sciences, Hatfield, PA). These PDMS stamps were sealed to glass coverslips with features-side down. A solution of 1% Agarose (#MIDSCI500 – Midsci, St. Louis, MO) in distilled water (C = 10 mg/mL) was heated to 100 °C until the solution became crystal clear. Subsequently, 600 mL of this 1% agarose solution was mixed with 400 mL of 100% ethanol, and a drop of this hot agarose/ethanol solution was pipetted against the side of the PDMS stamp to perfuse through the gaps formed between the silicon mold and the coverslip. After several hours, the PDMS stamp was carefully removed from the coverslip with fine-tipped forceps. However, the agarose layers tend to detach from the glass coverslip in long-term cell culture conditions (37 °C, 5% CO_2_). To solve this issue, we precoated the glass coverslips with an ultra-thin layer of polystyrene dissolved in chloroform (0.2 mg mL^−1^). A small drop of this mixture was added onto each coverslip, which was then spin-coated (∼100 rpm for 1-2 seconds followed by ∼3,000 rpm for 4-6 second) to spread the polystyrene uniformly. Finally, the coverslips were exposed to UV light in a tissue culture hood for 1 hour to graft the polystyrene layer prior to the addition of the agarose/ethanol solution. To setup an experiment, the coverslips holding the polystyrene/agarose layers were incubated with fibronectin in PBS (50 mg mL^−1^) for 1 h at 37 °C, rinsed twice with PBS and dried several minutes before cells were seeded on the coverslips in culture media. After 48 hours of incubation, the coverslips with the cells were placed in a 37 °C / 5% CO_2_ chamber for live imaging. The samples were imaged using a Leica SP8 inverted confocal microscope, equipped with a 40x oil objective. A z-stack of 8 images (1,024 x 1,024 pixels, z-step size 2.5 µm) was recorded every 10 min for 24 h. Fluorescent signals (488 and 561 nm) were collected sequentially to avoid any bleed-through using a 100 Hz scan rate. ImageJ (NIH, 64-bit Java 1.8.0_172) was used to produce a maximum intensity projection of each z-stack. All the resulting 2D images were assembled into time-lapse videos.

### CRISPR knockout

All single guide RNAs (sgRNAs) used in this study were designed by using the online software (MIT CRISPR Design Tool: http://crispr.mit.edu). The sequences of sgRNAs are included in Supplementary Table 3. sgRNAs were cloned into the pSpCas9(BB)-2A-eGFP (Addgene, PX458) plasmid by ligating annealed oligonucleotides with BbsI-digested vector. 4 μg plasmid carrying the specific sgRNA was then transfected into 1 × 10^6^–2 × 10^6^ dissociated mEpiSCs using NEPA21 electroporator (Nepa Gene). For generating large genomic fragment deletion at specific gene locus, two sgRNA plasmids were transfected simultaneously. eGFP^+^ cells were collected by FACS at 48 h after transfection and replated. Single clones were picked and expanded. Homozygous knockout clones were confirmed by genomic PCR, Sanger sequencing and western blotting.

### Plasmids

pCAG-IP-mKO and pCAG-IP-eGFP plasmids were gifts from T. Hishida. piggyBac-eGFP-H2B and piggyBac transposase plasmids were gifts from Aryeh Warmflash lab. piggyBac-mScarlet-H2B plasmid was generated by cloning mScarlet coding sequence and replaced eGFP coding sequence in piggyBac-eGFP-H2B. pSpCas9(BB)-2A-eGFP (PX458) plasmid was purchased from Addgene (plasmid 48138).

### Mouse embryo collection and electroporation

Homozygous *Mavs* knockout or WT C57BL/6 female mice (4–11 weeks old) were superovulated by intraperitoneal (IP) injection with 7.5 IU of PMSG (Prospec), followed by IP injection with 7.5 IU of hCG (Sigma-Aldrich) 48 h later. After mating with homozygous *Mavs* knockout or WT C57BL/6 male mice, respectively, embryos were harvested at E2.5 [the presence of a virginal plug was defined as embryonic day 0.5 (E0.5)] in KSOM-Hepes^5^ by flushing oviducts and uterine horns. Embryos were cultured in the defined modified medium (mKSOMaa)^5^ until blastocyst stage. For generating *Mavs* knockout blastocysts by CRISPR/Cas9 gene editing, mouse zygotes were collected from superovulated WT BDF1 female mice (3–4 weeks old) crossed with WT BDF1 males at E0.5 and electroporated with RNP complex of *Mavs* targeting sgRNA (100 ng μL^-1^) and Cas9 protein (Takara Bio; 500 ng μL^-1^) using NEPA21 electroporator. The electroporated zygotes were cultured in mKSOMaa until blastocyst stage. Embryo culture was performed in a humidified atmosphere containing 5% (v/v) CO_2_ and 20% (v/v) O_2_ at 37 °C.

### Microinjection of human iPS cells to mouse blastocysts

Microinjection of human iPS cells into mouse blastocysts was performed as previously described^10^ with slight modifications. Briefly, the embryos having obvious blastocoel at E3.5 were defined as blastocysts and fully expanded blastocysts were used for human iPS cell injections. Single cell suspensions of hiPS cells were added to a 40 μL droplet of KSOM-Hepes containing the blastocysts to be injected. Individual cells were collected into a 20 μm internal diameter of micropipette. Ten cells were introduced into the blastocoel near the ICM. Groups of 10–12 blastocysts were manipulated simultaneously, and each session was limited to 30 min. After microinjection, the blastocysts were cultured in mKSOMaa for at least 1 h until the *ex vivo* embryo culture or embryo transfer. For *ex vivo* embryo culture, we followed the previous protocol^25^. In brief, injected chimeric blastocysts were placed in ibiTreat μ-plate 8 wells (ibidi GabH, 80826) containing IVC1 medium (Cell guidance systems, M11) and cultured for 3 days in a humidified atmosphere containing 5% (v/v) CO_2_ and 5% (v/v) O_2_ at 37 °C. IVC1 medium was replaced with equilibrated IVC2 medium (Cell guidance systems, M12) and cultured for additional 2 days. Embryos were imaged using a fluorescence microscope (Echo Laboratories).

### Mouse embryo transfer

As surrogates, ICR female mice (8 weeks old or older) in the proestrus were mated with vasectomized ICR male mice to induce pseudopregnancy. Embryo transfer to the surrogate at E2.5 performed surgically under anesthesia with ketamine/xylazine mixture (ARC UTSW). hiPSC-injected blastocysts were loaded to the pipette with air bubble and transferred to the uterine horn, which were preliminary punctured with a 27G needle connected to a 1.5 mL syringe. 12–28 blastocysts were transferred per surrogate. After the transfer, opioid analgesic (Buprenorphine HCL) was treated. Pregnant female mice were euthanized on E8.5-E9.5, embryos were dissected and used for downstream analysis.

### Genomic PCR

Genomic PCR was performed for detecting human-specific DNA in mouse embryos by DNA fingerprinting using primers for *TPA25-Alu*. Genomic DNA of E8.5-E9.5 mouse embryos and HFF-hiPS cells (used as a positive control) were extracted using DNeasy Blood and Tissue Kit (Qiagen) following the manufacturer’s instructions, and then diluted to 30 ng μL^-1^ as PCR templates. Genomic PCRs were performed using Hot Start Taq 2x Master Mix (NEB). *PTGER2* primers that bind to conserved sequences in both human and mouse were used as internal control. The PCR products were examined by 2% agarose gel electrophoresis. Primer sequences are provided in Supplementary Table 3.

### Fluorescence imaging and immunohistochemistry analysis of mouse embryos

Imaging of chimeric embryos was performed with a Zeiss Axio Zoom.V16 fluorescence stereo zoom microscope equipped with a Plan-Neofluar Z 1.0x/0.25 (FWD 56 mm) objective and Axiocam 503 monochromatic camera. GFP-positive embryos were embedded in O.C.T. compound (Fisher Healthcare) without fixation and frozen on dry ice. Sections (12 μm thick) were cut on a cryostat (Leica CM 1950), mounted on the glass slides (Denville Scientific), and stored at -80 °C for immunohistochemistry. All extra-sections and GFP-negative embryos were subjected to DNA extraction and genotyping for *Mavs*. For immunohistochemistry, sections were fixed in 4% paraformaldehyde (PFA) for 30 min at 4 °C, washed with PBS twice and permeabilized with 0.3% Triton X-100 (Fisher BioReagents) in PBS twice for 5 min at room temperature. After antigen retrieval with 10 mM Citrate (pH 6.0) containing 0.05% Tween 20 (Fisher BioReagents) for 10 min at 90-95 °C, sections were blocked with blocking buffer [5% (w/v) donkey serum (Sigma-Aldrich) and 0.3% (v/v) Triton X-100 in PBS] for 1 h. Then sections were incubated with the primary antibody (chicken anti-GFP, Supplementary Table 4) diluted in blocking buffer at 4 °C overnight, secondary antibody (Donkey anti-chicken IgY conjugated with FITC, Supplementary Table 4) in blocking buffer at room temperature for 1 h, and finally 5 ng/mL DAPI solution at room temperature for 10 min. Autofluorescence was silenced with TrueBlack Lipofuscin Quencher (Biotium, diluted 1:20 in 70% ethanol) and slides were mounted with PVA-DABCO. Samples were imaged using a confocal microscope (Zeiss LSM 700).

### Immunofluorescence

Cells grown on coverslips were fixed in 4% paraformaldehyde (PFA) for 10 min at room temperature, permeabilized with 0.1% Triton X-100 for 15 min, and blocked with 5% BSA and 0.1% Triton X-100 for 1 h. Incubation with primary antibodies (Supplementary Table 4) in 1% BSA and 0.1% Triton X-100 was performed overnight at 4 °C. Incubation with secondary antibodies (Supplementary Table 4) in 1% BSA and 0.1% Triton X-100 was performed for 1 h at room temperature. DAPI staining was applied for 10 min at room temperature. Coverslips were then mounted with PVA-DABCO on glass slides. The images of stained slides were taken by Revolve (ECHO) or Leica SP8 confocal microscope.

### Flow cytometry

Cells were dissociated using TrypLE and fixed in 4% PFA for 10 min at room temperature. Then cells were resuspended with PBS and subjected to flow cytometry analysis. Flow cytometry was performed using FACSCalibur system (BD Bioscience) and analyzed using FlowJo (10.8.1).

### Western blotting and Native gel electrophoresis

For western blotting, cells were lysed with RIPA Lysis Buffer (Thermo Fisher Scientific) with protease inhibitor on ice and centrifuged at 12,000 *g*. The supernatant was mixed with 4× SDS Loading Buffer (Bio-Rad) and boiled at 95°C for 10 min. Samples were loaded into 4-20% Tris-Glycine gel (Invitrogen Novex WedgeWell 4 to 20%, Tris-Glycine) and run at 150 V for 60 min. The gels were then transferred to PVDF membranes (Bio-Rad) and blocked with 5% milk in TBST. Membranes were incubated with the corresponding primary antibodies (Supplementary Table 4). Immunoreactive bands were visualized using HRP conjugated secondary antibodies (Supplementary Table 4) incubated with chemiluminescence substrate (Pierce ECL western substrate, Thermo Fisher Scientific) and exposed to X-ray film. For native gel electrophoresis, cells were lysed with Lysis Buffer (20 mM Tris-HCl, pH 7.5, 150 mM NaCl, 10% Glycerol, 1% NP-40, 5 mM Na_3_VO_4_ and protease inhibitor) on ice followed by 12,000 *g* centrifugation. The supernatant was mixed with 5× Native Loading Buffer (125 mM Tris, 0.96 M Glycine, 40% Glycerol, 0.005% Bromophenol Blue and 2% DOC-Na). The Cathode Buffer (25 mM Tris, 192 mM Glycine and 0.4% DOC-Na) and the Anode Buffer (25 mM Tris and 192 mM Glycine) were applied. 10% Tris-Glycine gel (Invitrogen Novex 10% Tris-Glycine gel) was pre-run at 200V for 70 min in the cold room until current dropped to 10 mA/Gel. The samples were loaded and run at 200 V for 60 min in the cold room. The gels were then transferred, blocked with 5% milk in TBST and blotted with anti-IRF3 antibody (Supplementary Table 4).

### RNA isolation and quantitative RT–PCR analysis

Both co-cultured and separately cultured cells were sorted out and collected based on fluorescent labelling using FACS at each of the indicated time points. Total RNAs were extracted using RNeasy Mini Kit (Qiagen). cDNA was synthesized using iScript™ Reverse Transcription Supermix (Bio-Rad). Quantitative reverse transcription PCR (qRT–PCR) was carried out using SYBR Green Master Mix (Qiagen) on CFX384 system (Bio-Rad). Reactions were run in triplicate and expression level of each gene was normalized to the geometric mean of Gapdh as a housekeeping gene and analysed by using the ΔΔCt method by Bio-Rad Maestro 1.0. The qRT– PCR primer sequences of each gene are listed in Supplementary Table 3.

### Bulk RNA-seq quality control and data analysis

RNA-seq reads were first filtered by SOAPnuke (Version: 1.5.6)^42^ with parameters “-l 15 -q 0.2 -n 0.05 -Q 2”. The high-quality reads were mapped to the mouse genome (2020-A, mm10) and the human genome (2020-A, GRCh38) downloaded from the 10X Genomics website using HISAT2 (Version: 2.1.0)^43^ with parameters “-k 1 -p 2 -S --novel-splicesite-outfile -q --no-unal --dta --un-conc-gz”. SAMtools (Version: 1.5)^44^ was used to sort the BAM files produced by HISAT2. The expression levels of annotated genes were calculated by transcript per million (TPM) using StringTie (Version: 2.1.4)^45^ with parameters “-t -C -e -B -A -p 2”. Fold change was calculated by TPM+1 to avoid infinity. A fold change >= 2 or <= 0.5, a *P* value and an adjusted *P* value calculated by DESeq2^46^ less than 0.05 and 0.1, were used as cutoffs to define differentially expressed genes. KEGG pathway enrichment analyses of up-regulated mouse and human genes were performed by DAVID^47^ (Version: 2021 Update) with mouse (*Mus musculus*) and human (*Homo sapiens*) as the backgrounds. Default thresholds (Count > 2, EASE < 0.1) were used to define enriched KEGG pathways.

### scRNA-seq quality control and data analysis

scRNA-seq reads were processed by Cell Ranger (Version: 5.0.1) using parameters “--localcores=24 --localmem=400 --transcriptome --fastqs” with the mouse genome annotation (2020-A, mm10) and the human genome annotation (2020-A, GRCh38). Multigenome analysis using the human-mouse genome (GRCh38-and-mm10-2020-A) of Cell Ranger was performed to distinguish mouse cells and human cells from cross-species doublets. scRNA-seq analysis was performed using Seurat (Version: 4.0.4)^48^. DoubletFinder (Version: 2.0.3)^49^ was used to remove intra-species doublets with parameters “pN = 0.25, nExp =round(0.05*length(colnames(obj)))”. Clusters of mouse embryonic fibroblasts (MEFs) were removed based on the following features: (1) they were present in all samples; (2) they were mouse cells; (3) they expressed marker genes of MEFs; and (4) they did not express marker genes of mEpiSCs. For mouse cells, cells with less than 500 genes or UMI less than 1,000 or mitochondrial ratio greater than 10% were removed. Finally, a total of 16,225 mouse cells were kept for further analysis. Cell clustering and Uniform Manifold Approximation and Projection (UMAP) visualization were performed using the “FindClusters” and “RunUMAP” functions of Seurat, respectively.

### Identification of mouse-/human-specific and eGFP-/hKO-specific reads

To identify species-or fluorescent gene-specific reads, the high-quality bulk RNA-seq of each sample and scRNA-seq reads of each cell were mapped to both mouse and human genomes, or eGFP and hKO sequences, respectively. Reads that could be mapped to both genomes or both fluorescent gene sequences were removed. This process was repeated many times until there were only species-or fluorescent gene-specific reads left, which were used to calculate mouse/human genome mapping rates, measure eGFP/hKO expression levels, and detect eGFP^+^/hKO^+^ cells. Reads per million (RPM) was used to calculate normalized expression levels of human transcripts and eGFP/hKO. eGFP^+^/hKO^+^ cells were defined as cells with eGFP-/hKO-specific reads >=1.

### Retrotransposable element analysis

RNA sequencing fastq files were preprocessed with Cutadapt v. 1.18 and Prinseq v. 0.20.4 to remove residual adapter sequences, low quality reads and bases^50, 51^. Remaining reads were aligned with STAR v. 2.6.0 (--twopassMode Basic --outFilterMultimapScoreRange 2 -- winAnchorMultimapNmax 1000 --outFilterMultimapNmax 10000 --outFilterMismatchNmax 20 --scoreDelOpen -1 --scoreDelBase -1 --scoreInsOpen -1 --scoreInsBase -1) to both the hg38 assembly of the human genome and the mm10 assembly of the mouse genome using the UCSC transcriptome annotation^52^. Picard v. 2.18.9 (http://broadinstitute.github.io/picard/) and SAMtools v. 1.9 were used to identify and filter PCR duplicates^44^. Read counting was performed with HTseq v. 0.11.0 using only unique reads aligning to UCSC annotated gene transcripts^53^. The resulting gene counts file was used for Deseq2 v 2.11.40.6 TMM normalization and differential gene expression analysis^46^. Normalized stranded bigwig files for visualization were generated using the Deeptools v 3.0.1 BamCoverage tool (--binSize 5 --normalizeUsing CPM --minMappingQuality 255 --filterRNAstrand forward/reverse --samFlagExclude 1024)^54^. Retroelement expression levels were determined using a customized pipeline^55^. A general transfer format (GTF) file containing RepeatMasker retroelement annotations was generated for the hg38 human and mm10 mouse genome assemblies^56^. This annotation file contains the genomic coordinates, strand, conservation scores relative to consensus sequence, and relational information for RepeatMasker each annotated repetitive element (element name, repeat family, and repeat class). We further identified and denoted the most highly conserved, full length retroelement copies present in these genomes. Reads aligning to individual repetitive elements, reads aligning to specific families of repeat elements, and reads aligning to repetitive sequences classes were counted using HTseq^53^. Reads multiply mapped to more than one individual copy of an element, element within a given family or class of elements were counted only once when calculating read counts at the element, class or family levels. Repetitive sequence read counts were normalized using the Deseq2 scale factors generated above. As a quality control metric, reads derived from the opposite strand from the retroelement-encoding strand were also counted.

### Statistics

Data were presented as mean ± s.e.m. from at least three independent experiments. Differences between groups were evaluated by Student’s t-test (two-sided) or one-way ANOVA with Dunnett’s multiple comparison, and considered to be statistically significant if P < 0.05. Graphic analyses were done using GraphPad Prism version 9.0 (GraphPad Software). Statistical analyses were done using the Microsoft Excel (Microsoft 365).

### Reporting summary

Further information on research design is available in the Nature Research Reporting Summary linked to this paper.

### Data availability

The RNA-seq datasets generated in this study have been deposited in the CNGB Sequence Archive (CNSA; https://db.cngb.org/cnsa/)^57^ of the China National GeneBank DataBase (CNGBdb)^58^. They will be made publicly accessible upon publication.

---

**Extended Data Fig. 1.**
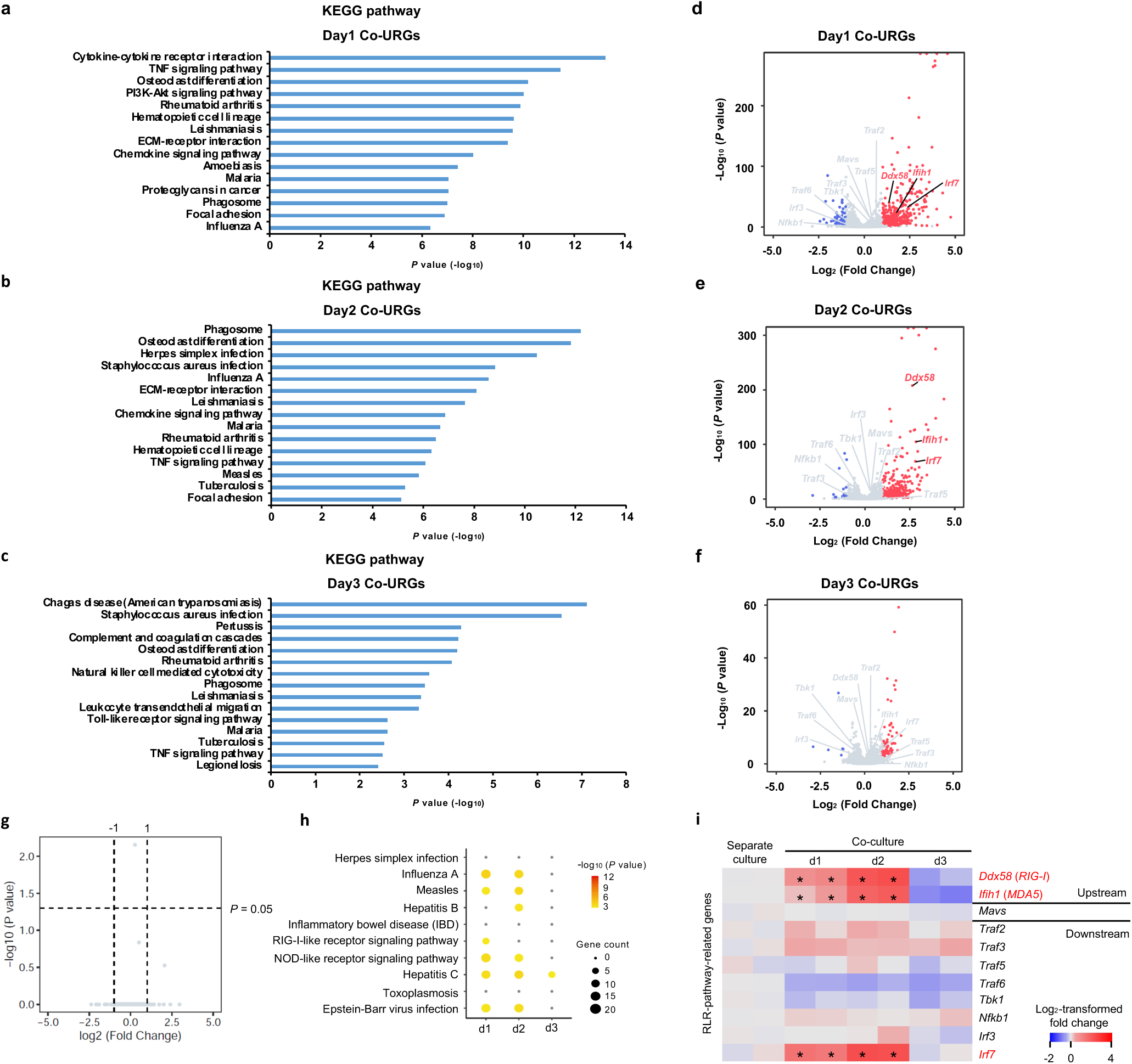
| Comparative RNA-seq analysis between co-cultured and separately cultured mEpiSCs. **a-c,** KEGG pathways enriched on days 1 (**a**), 2 (**b**), and 3 (**c**) co-URGs in mEpiSCs. Top 15 (with the lowest *P* values) KEGG pathways are shown. **d-f,** Volcano plots showing significantly upregulated (red) and downregulated (blue) genes in co-cultured versus separately cultured mEpiSCs on days 1 (**d**), 2 (**e**) and 3 (**f**). RLR-pathway-related genes are highlighted. **g,** Volcano plot showing no differentially expressed genes in transwell co-cultured versus separately cultured mEpiSCs. **h,** KEGG pathway analysis of upregulated genes in mEpiSCs co-cultured with *P65*^KO^ hiPSCs (versus separate cultures). The color of each dot represents log_10_-transformed *P* value of each term, while the dot size represents the number of co-URGs in each term. Terms that are not significantly enriched are shown in gray. **i,** Heatmap showing fold changes of RLR-pathway-related genes in mEpiSCs co-cultured with *P65*^KO^ hiPSCs (versus separate cultures). Blue and red represent log_2_-transformed fold changes < 0 and > 0, respectively. Asterisks indicate statistically differentially expressed.

**Extended Data Fig. 2.**
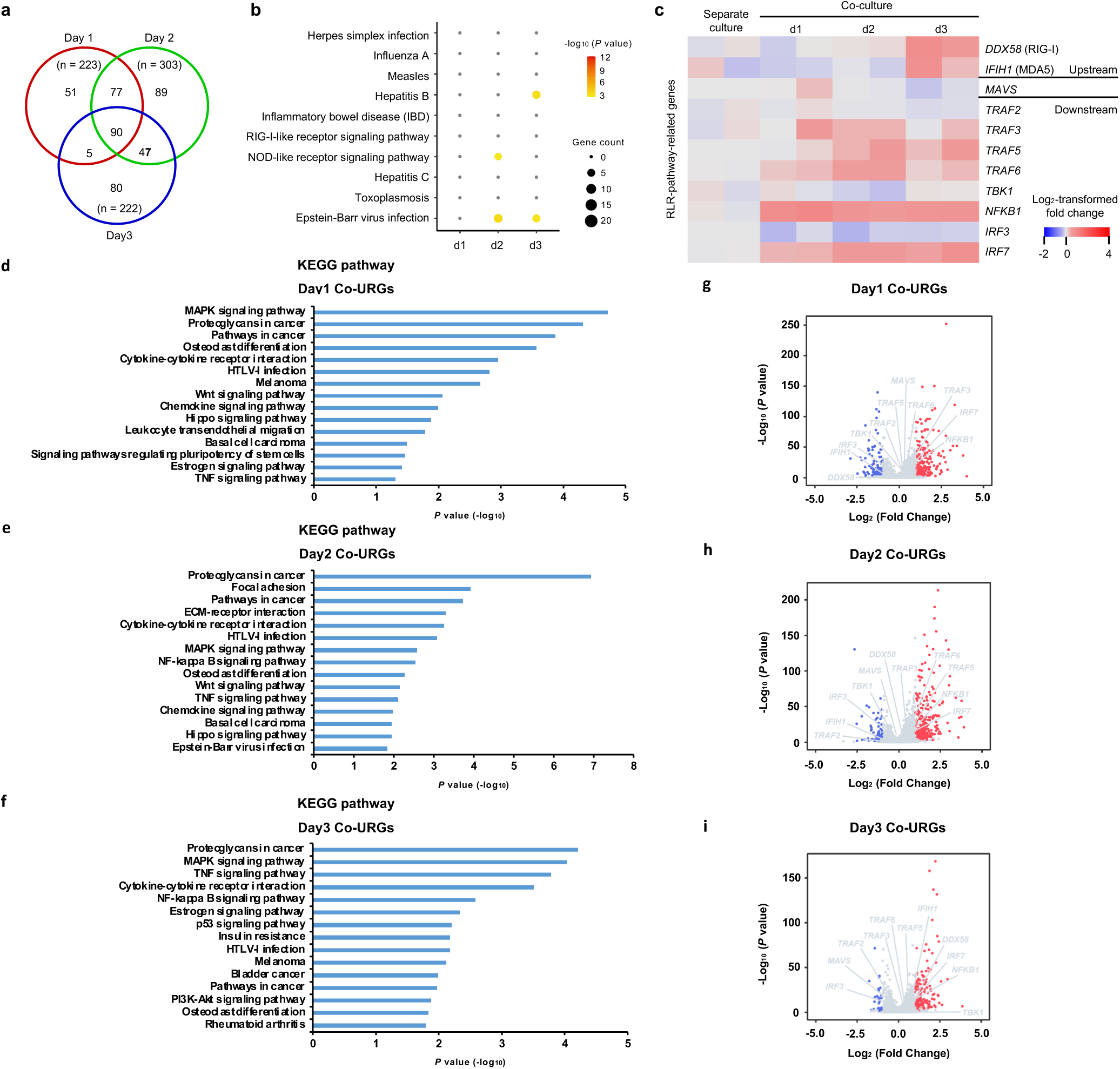
| Comparative RNA-seq analysis between co-cultured and separately cultured H9 hES cells. **a,** Venn diagram showing the numbers of co-URGs in H9 hES cells. **b,** KEGG pathway analysis of co-URGs in H9 hES cells. The color of each dot represents log_10_-transformed *P* value of each term, while the dot size represents the number of co-URGs in each term. Terms that are not significantly enriched are shown in gray. **c,** Heatmap showing fold changes of RLR-pathway-related genes in co-cultured versus separately cultured H9 hES cells. Blue and red represent log_2_-transformed fold changes > 0 and < 0, respectively. **d-f,** KEGG pathways enriched on days 1 (**d**), 2 (**e**), and 3 (**f**) co-URGs in H9 hES cells. Top 15 (with the lowest *P* values) KEGG pathways are shown. **g-i,** Volcano plots showing significantly upregulated (red) and downregulated (blue) genes in co-cultured versus separately cultured H9 hES cells on days 1 (**g**), 2 (**h**) and 3 (**i**). RLR-pathway-related genes are highlighted.

**Extended Data Fig. 3.**
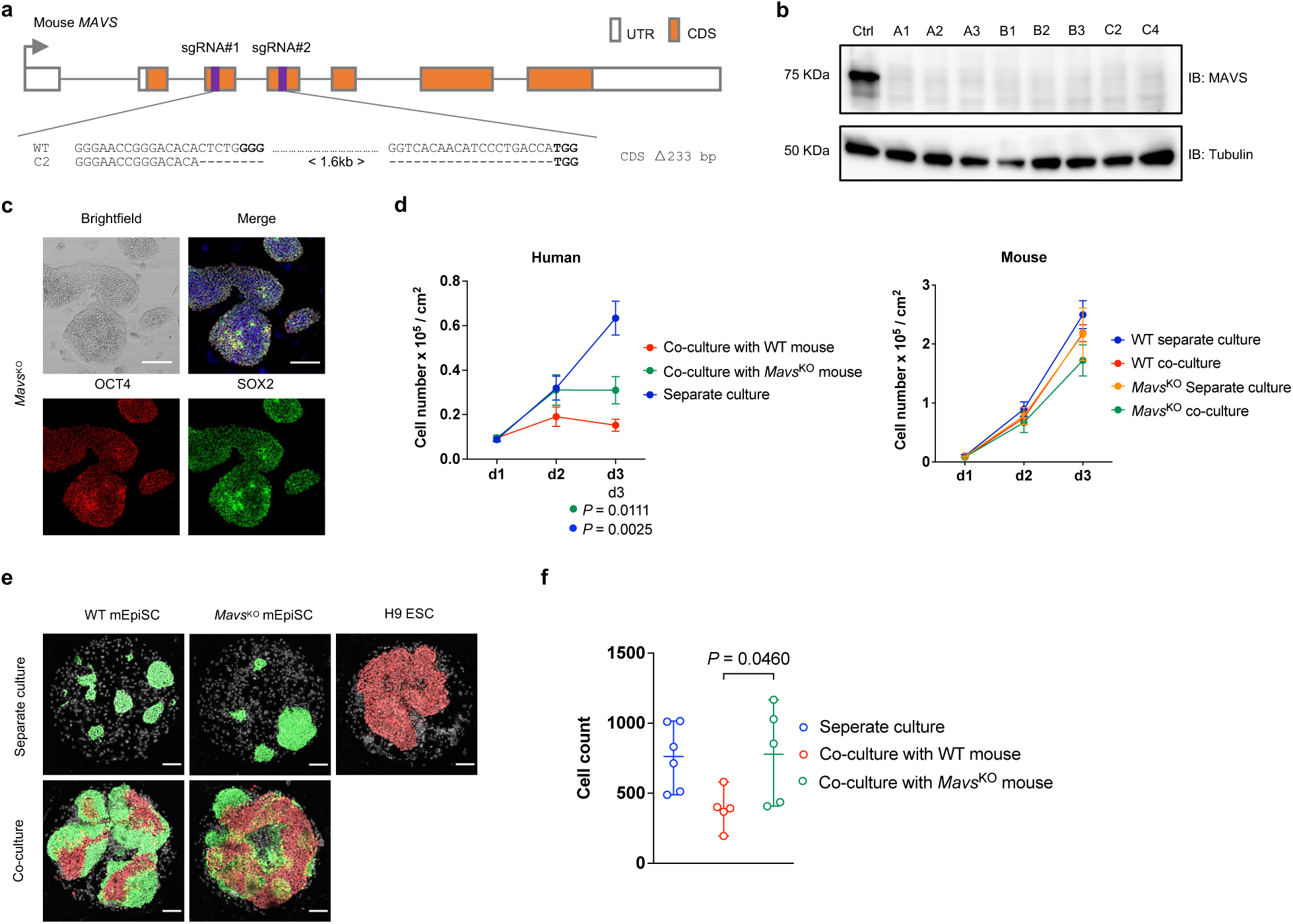
| Genetic inactivation of *Mavs* in mEpiSCs compromises winner cell competitiveness. **a**, Schematic of mouse *Mavs* gene targeting with two sgRNAs and homozygous 1.6-kb deletion (233-bp out-of-frame deletion in CDS) in mEpiSC clone C2. Bold, PAM sequence. **b,** Western blot confirmed the lack of MAVS protein expression in several independent *Mavs*^KO^ mEpiSC clones. Tubulin was used as a loading control. **c,** Representative brightfield and immunofluorescence images showing long-term cultured *Mavs*^KO^ mEpiSCs maintained stable colony morphology and expressed core (OCT4, red; SOX2, green) pluripotency markers. Blue, DAPI. Scale bars, 100 μm. **d,** Growth dynamics of human-mouse 3-day-co-culture assay. Left, growth curves of H9 hES cells in co-culture with WT mEpiSCs (red), co-culture with *Mavs*^KO^ mEpiSCs (green) and separate culture (blue). Right, growth curves of WT mEpiSCs in co-culture (red) with H9 hES cells and separate culture (blue), *Mavs*^KO^ mEpiSCs in co-culture (green) with H9 hES cells and separate culture (orange). n = 4, biological replicates. Data are mean ± s.e.m. *P* values determined by one-way ANOVA with Dunnett’s multiple comparison. **e,** Representative fluorescence images of day-3 co-cultured and separately cultured WT mEpiSCs (green), *Mavs*^KO^ mEpiSCs (green) and H9 hES cells (red) in MEF-coated micropattern wells. White, DAPI. Scale bar, 100 μm. **f,** Dot-plot showing numbers of live H9 hES cells in separate culture (n = 6), co-culture with WT mEpiSCs (n = 5) and co-culture with *Mavs*^KO^ mEpiSCs (n = 5) on day 3 in micropatterned wells. n, biological replicates.

**Extended Data Fig. 4.**
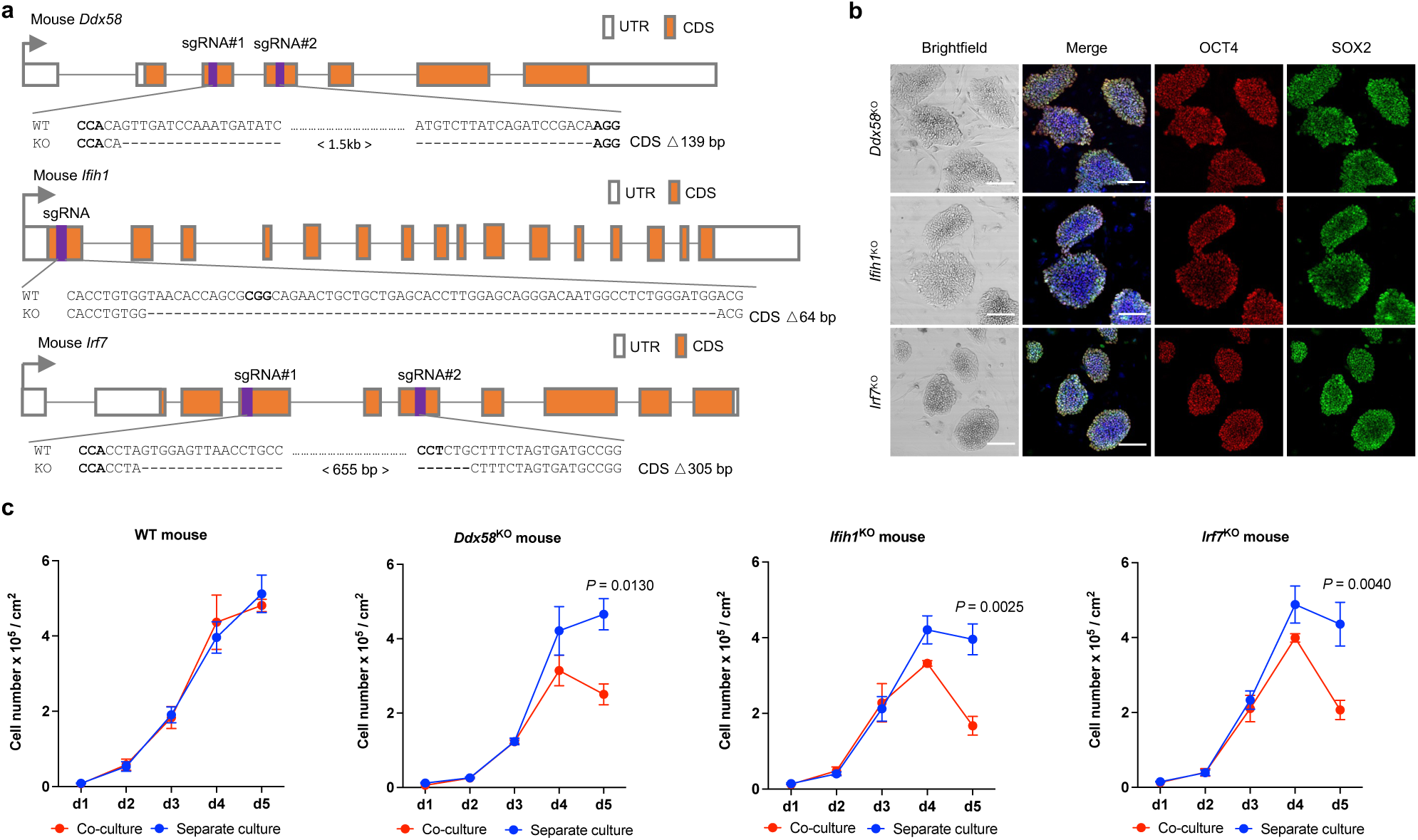
| Genetic inactivation of *Ddx58*, *Ifih1* and *Irf7* in mEpiSCs compromise winner cell viability during human-mouse PSC co-culture. **a**, Upper, schematic of mouse *Ddx58* gene targeting with two sgRNAs and homozygous 1.5-kb deletion (139-bp out-of-frame deletion in CDS) in *Ddx58*^KO^ clone. Middle, schematic of mouse *Ifih1* gene targeting with one sgRNA and homozygous 64-bp out-of-frame deletion in *Ifih1*^KO^ clone. Lower, schematic of mouse *Irf7* gene targeting with two sgRNAs and homozygous 655-bp deletion (305-bp out-of-frame deletion in CDS) in *Irf7*^KO^ clone. Bold, PAM sequence. b, Representative brightfield and immunofluorescence images showing long-term cultured *Ddx58*^KO^, *Ifih1*^KO^ and *Irf7*^KO^ mEpiSCs maintained stable colony morphology and expressed core (OCT4, red; SOX2, green) pluripotency markers. Blue, DAPI. Scale bars, 100 μm. c, Growth curves of co-cultured and separately cultured WT, *Ddx58*^KO^, *Ifih1*^KO^ and *Irf7*^KO^ mEpiSCs. n = 6 (WT), n = 3 (*Ddx58*^KO^), n = 3 (*Ifih1*^KO^) and n = 3 (*Irf7*^KO^) independent culture experiments. Data are mean ± s.e.m. *P* values determined by unpaired two-tailed t-test.

**Extended Data Fig. 5.**
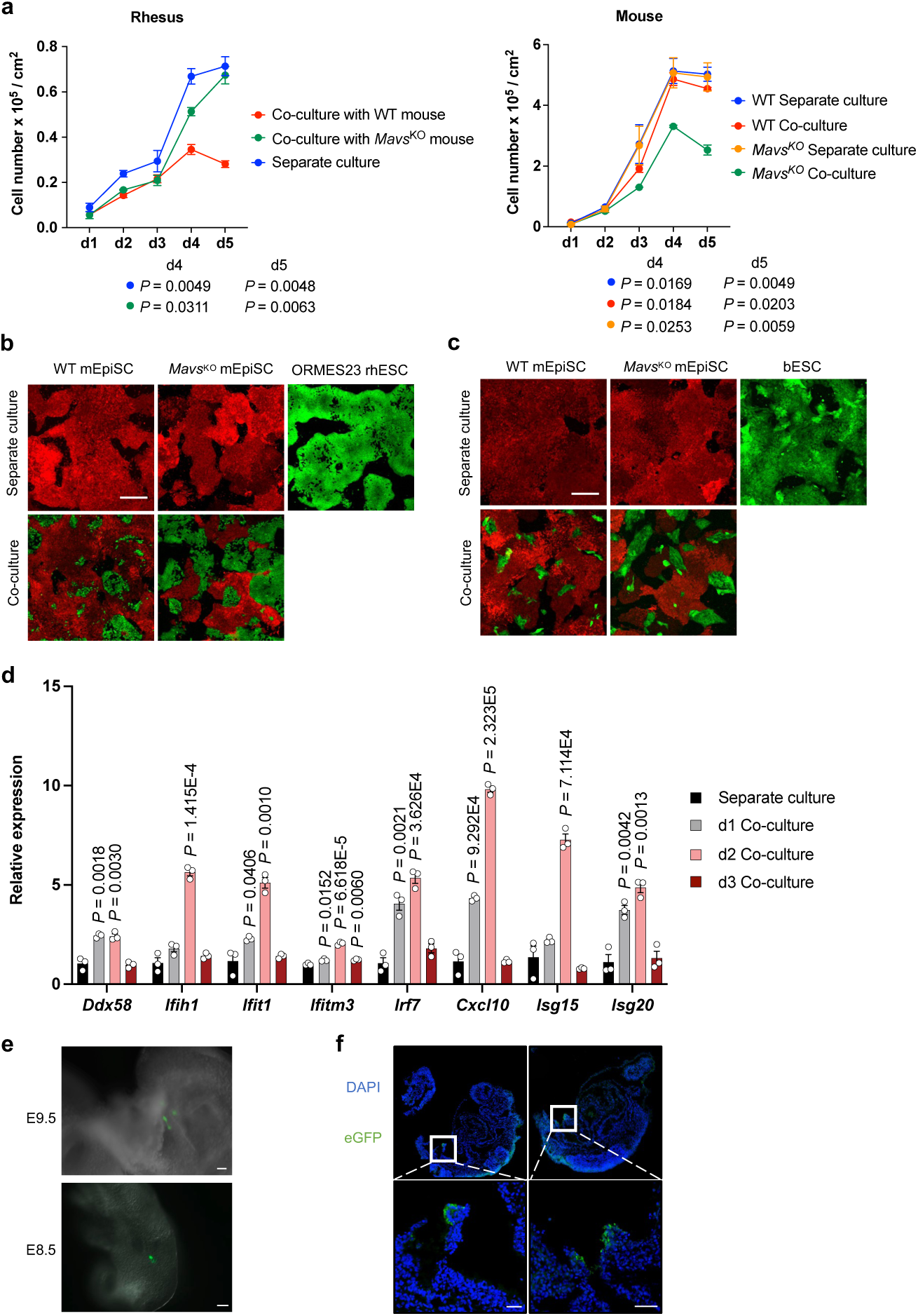
| Conserved role of *Mavs* during competitive interactions of mEpiSCs with primed PSCs from other species. **a,** Left, growth curves of rhesus ES cells (ORMES23) in co-culture with WT mEpiSCs (red), co-culture with *Mavs*^KO^ mEpiSCs (green) and separate culture (blue). Right, growth curves of WT mEpiSCs in co-culture (red) with rhesus ES cells (ORMES23) and separate culture (blue), *Mavs*^KO^ mEpiSCs in co-culture (green) with rhesus ES cells (ORMES23) and separate culture (orange). n =3, biological replicates. All data are mean ± s.e.m. *P* values determined by one-way ANOVA with Dunnett’s multiple comparison. **b,** Representative fluorescence images of day-4 co-cultured and separately cultured WT mEpiSCs (red), *Mavs*^KO^ mEpiSCs (red) and rhesus ES cells (ORMES23) (green). Scale bar, 200 μm. **c,** Representative fluorescence images of day-4 co-cultured and separately cultured WT mEpiSCs (red), *Mavs*^KO^ mEpiSCs (red) and bovine ES cells (green). Scale bar, 200 μm. **d,** RT-qPCR analysis of relative expression levels of RLR-pathway-related genes in days 1, 2 and 3 co-cultured (with bovine ES cells) *Mavs*^KO^ mEpiSCs compared to separately cultured *Mavs*^KO^ mEpiSCs. n = 3, biological replicates. Data are mean ± s.e.m. *P* values determined by unpaired two-tailed t-test. **e,** Representative fluorescence images of E8.5-9.5 *Mavs*^KO^ mouse embryos showing eGFP signal after blastocyst injection of eGFP-labeled hiPSCs and embryo transfer. Scale bars, 50 μm. **f,** Representative immunofluorescence images showing contribution of eGFP-labeled hiPSCs in E8.5-9.5 mouse embryos. Embryo sections were stained with antibody against eGFP and DAPI. Scale bars, 50 μm.

**Extended Data Fig. 6.**
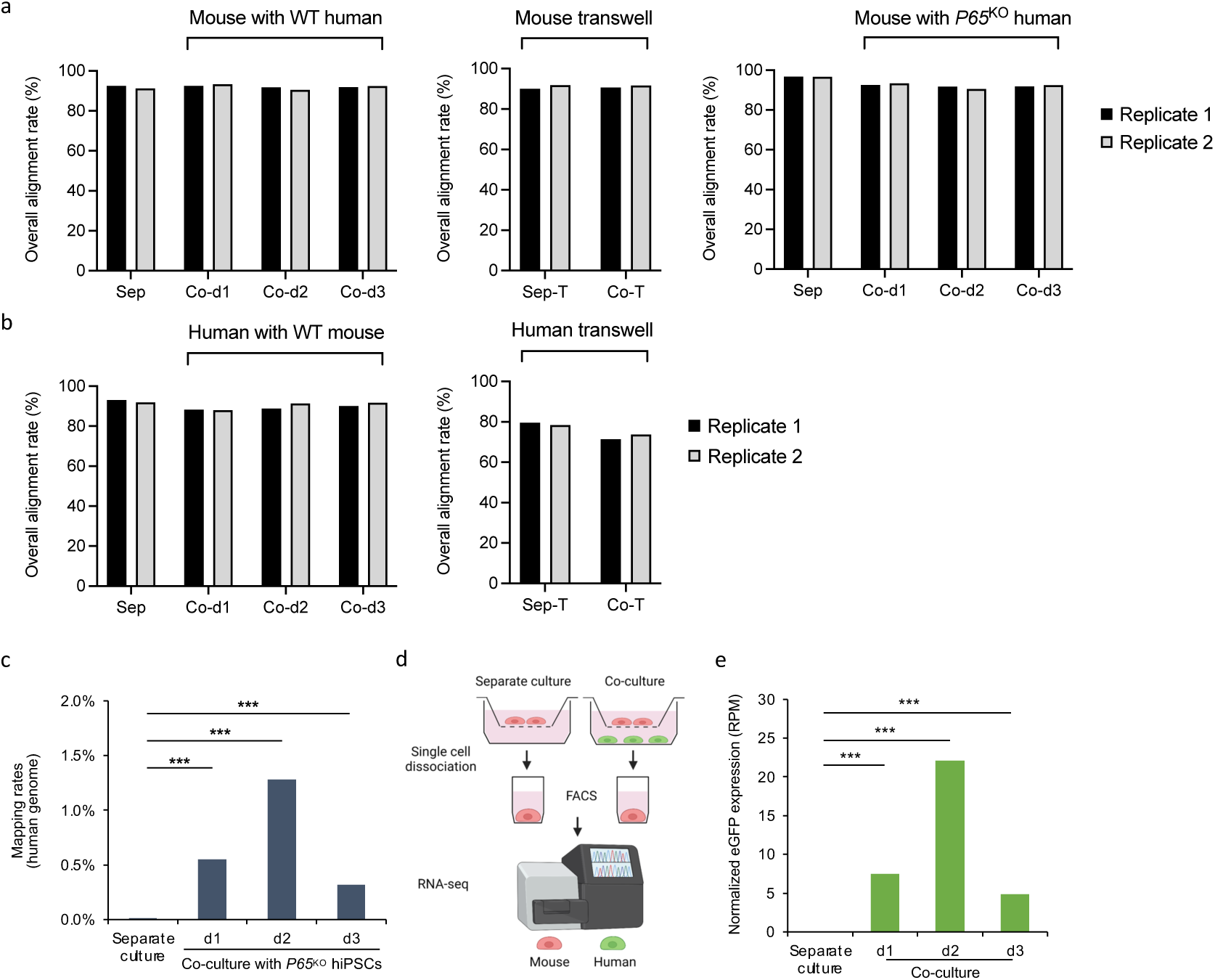
| Presence of human transcripts in co-cultured mEpiSCs revealed by bulk RNA sequencing. **a,** Statistics of bulk RNA-seq reads in mEpiSCs that could map to mouse genome. **b,** Statistics of bulk RNA-seq reads in H9 hES cells that could map to human genome. For **a** and **b**, Sep, Co-d1, Co-d2, Co-d3, Sep-T and Co-T represent separate culture, co-culture on days 1, 2, 3, separate culture in transwell and co-culture in transwell, respectively. **c,** Mapping rates of human genome in mEpiSCs co-cultured with *P65*^KO^ hiPSCs determined by bulk RNA-seq. Asterisks indicate statistically significant differences: (***) *P* < 0.001; Fisher’s exact test. **d,** Schematic of human and mouse PSC co-culture in transwells and the following RNA-seq experimental setup. **e,** Normalized eGFP expression in mEpiSCs determined by bulk RNA-seq. Asterisks indicate statistically significant differences: (***) *P* < 0.001; Fisher’s exact test.

**Extended Data Fig. 7.**
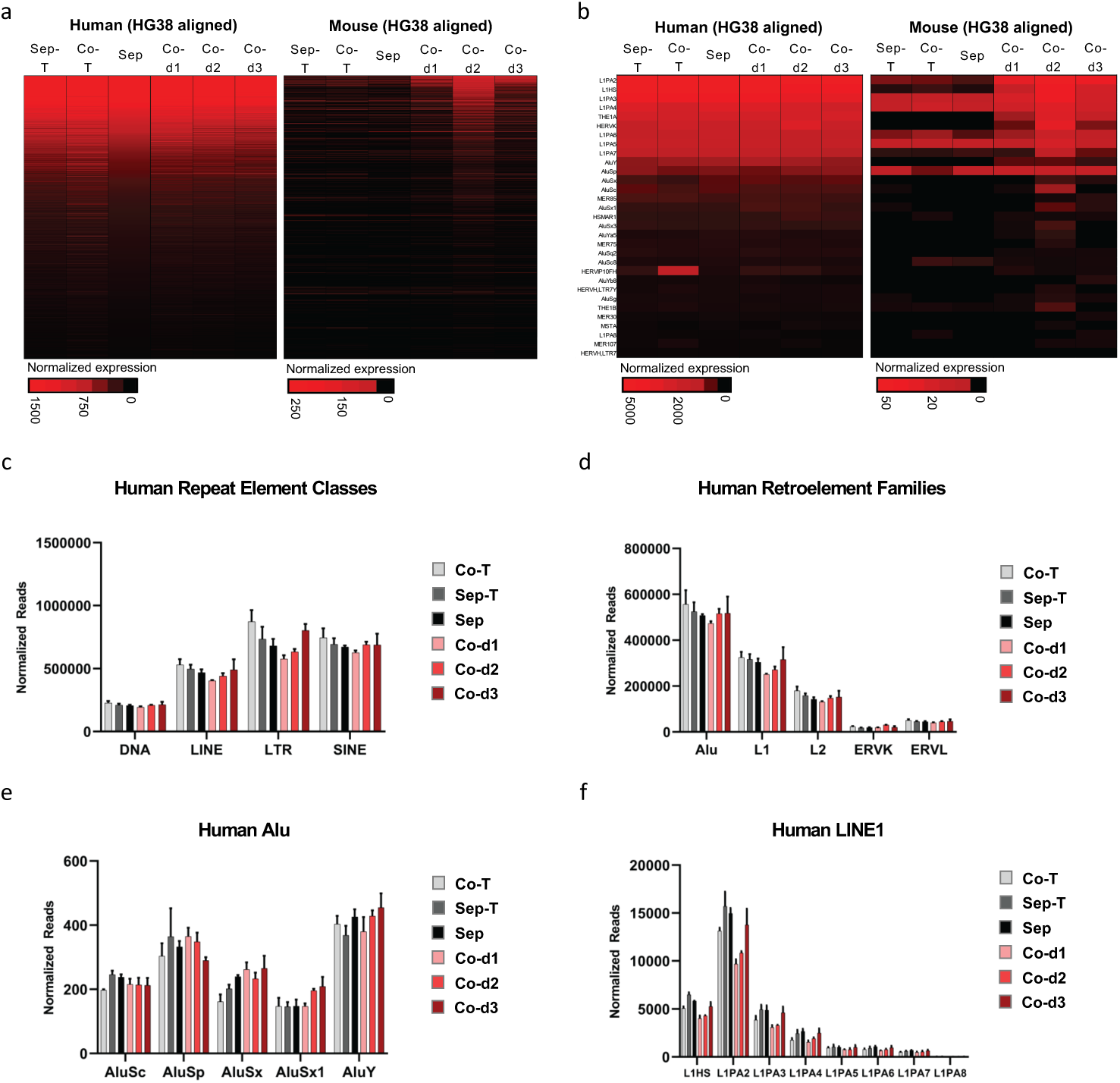
| Presence of RNAs encoding human repeat elements in co-cultured mEpiSCs revealed by bulk RNA sequencing. **a**, Normalized abundance of human repeat elements in H9 hES cells (left) or mEpiSCs (right) aligned to the human genome (assembly HG38). Shown are repeat elements expressed in separately cultured H9 hES cells (Normalized Expression >20) and not detected in separately cultured mEpiSCs (Normalized Expression <20). Only reads uniquely assignable to a single repeat element type are shown. Heat maps were sorted on expression from high to low in H9 hES cells. **b**, Normalized abundance of near-consensus human repeat elements in H9 hES cells (left) or mEpiSCs (right). Shown are least divergent repeat elements (full-length relative to consensus, <10 mismatches or indels) that were expressed in separately cultured H9 hES cells (Normalized Expression >20) and were not detected in separately cultured mEpiSCs (Normalized Expression <20). Heat maps were sorted on expression from high to low in H9 hES cells. **c-f**, Expression of human repeat element classes (**c**), retroelement families (**d**), Alu (**e**) and evolutionarily young LINE1 (**f**) elements in H9 hES cells under different culture conditions. Sep, Co-d1, Co-d2, Co-d3, Sep-T and Co-T represent separate culture, co-culture on days 1, 2, 3, separate culture in transwell and co-culture in transwell, respectively.

**Extended Data Fig. 8.**
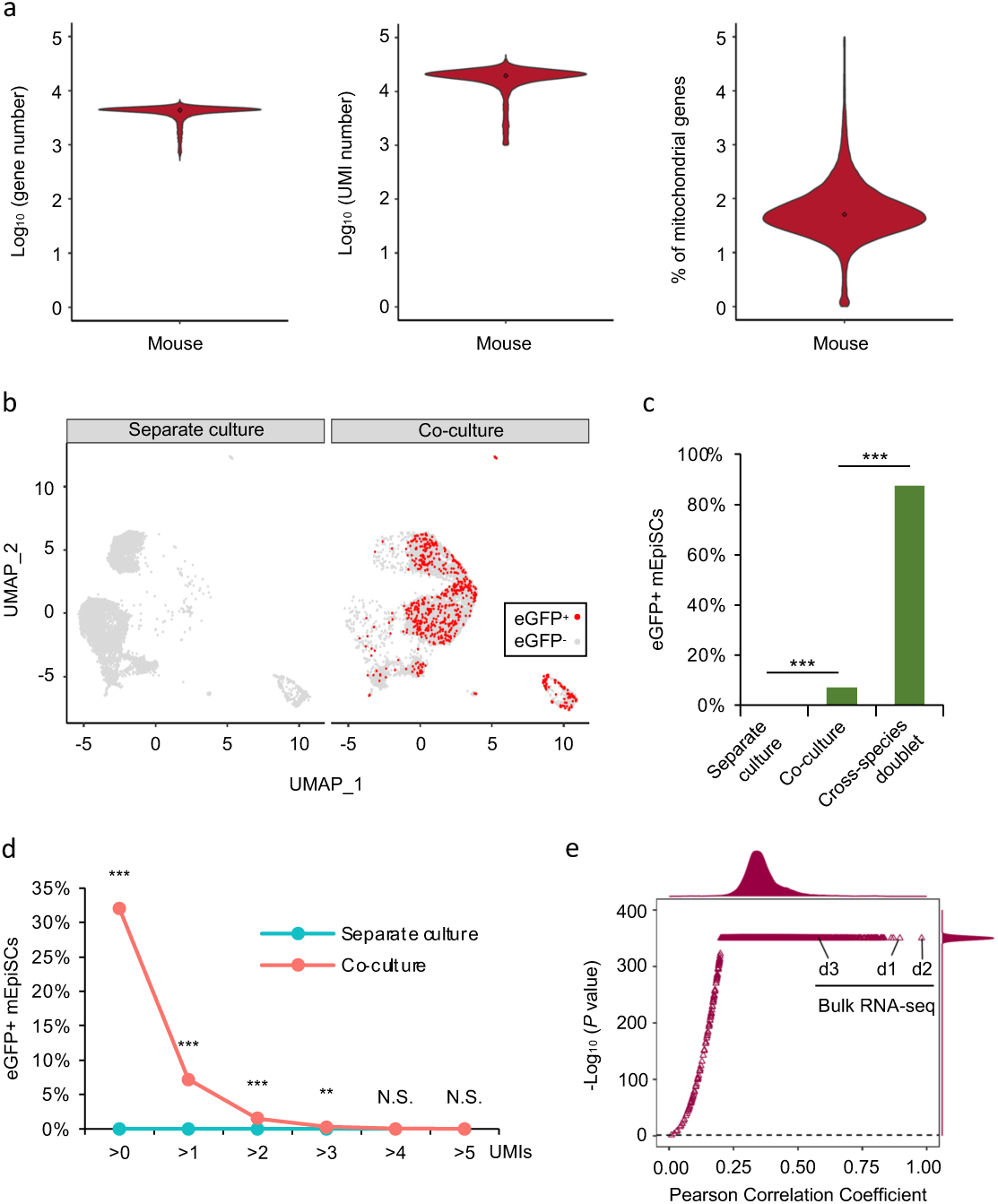
| Presence of human transcripts in co-cultured mEpiSCs revealed by single cell RNA sequencing. **a,** Violin plots showing the number of genes (left), UMIs (middle) and the percentage of mitochondria genes (right) per cell. **b,** UMAP visualization showing eGFP^+^ (red) and eGFP^-^ (gray) mEpiSCs in separate culture (left) and co-culture (right). **c,** Proportion of eGFP^+^ mEpiSCs determined by scRNA-seq. Asterisks indicate statistically significant differences: (***) *P* < 0.001; Fisher’s exact test. Cross-species doublets served as a negative control. **d,** Cumulative distribution showing proportion of eGFP^+^ mEpiSCs with different thresholds. Asterisks indicate statistically significant differences: (**) *P* < 0.01; (***) *P* < 0.001; Fisher’s exact test; (N.S.) not significant. **e,** Scatterplot showing global expression correlation of human genes detected in mEpiSCs and H9 hES cells determined by bulk RNA-seq and scRNA-seq. X-axis represents Pearson Correlation Coefficients whereas Y-axis represents *P* values. Triangles represent bulk RNA-seq datasets (indicated in the plot) or single-cell datasets. All datasets are summarized and shown as density plots along each axis.

