## Supplementary Information for "RNA Sensing and Innate Immunity Constitutes a Barrier for Interspecies Chimerism"

**Supplementary Table 1**

KEGG enrichment analyses of upregulated genes in co-cultured versus separately cultured mEpiSCs. *P* values determined by hypergeometric test.

**Supplementary Table 2**

KEGG enrichment analyses of upregulated genes in co-cultured versus separately cultured H9 hES cells. *P* values determined by hypergeometric test.

**Supplementary Table 3**

Sequence information of sgRNAs, sequencing primers, qRT-PCR primers and genomic PCR primers used in this study.

**Supplementary Table 4**

Information of primary and secondary antibodies used in this study.

**Supplementary Video 1**

mEpiSCs (left: WT; right: *Mavs*^KO^) and H9 hES cells co-culture (micropatterned cover slide). Time-lapse imaging of co-cultured H9 hES cells (green) and mEpiSCs (red) on micropatterned cover slide between days 2 and 3.

**Supplementary Video 2**

mEpiSCs (left: WT; right: *Mavs*^KO^) and bovine ES cells co-culture (micropatterned cover slide). Time-lapse imaging of co-cultured bovine ES cells (green) and mEpiSCs (red) on micropatterned cover slide between days 2 and 3.
